# Estimating daily intakes of manganese due to breast milk, infant formulas, or young child nutritional beverages in the United States and France: Comparison to sufficiency and toxicity thresholds

**DOI:** 10.1101/2020.06.09.142612

**Authors:** Erika J. Mitchell, Seth H. Frisbie, Stéphane Roudeau, Asuncion Carmona, Richard Ortega

## Abstract

**Background:** Although manganese (Mn) is an essential nutrient, recent research has revealed that excess Mn in early childhood may have adverse effects on neurodevelopment.

**Methods:** We estimated daily total Mn intake due to breast milk at average body weights by reviewing reported concentrations of breast milk Mn and measurements of body weight and breast milk intake at 3 weeks, 4.25 months, 7 months, and 18 months. We compared these figures to the Mn content measured in 44 infant, follow-up, and toddler formulas purchased in the United States and France. We calculated Mn content of formula products made with ultra-trace elemental analysis grade water (0 µg Mn/L) and with water containing 250 µg Mn/L, a concentration which is relatively high but less than the World Health Organization Health-based value of 400 µg Mn/L or the United States Environmental Protection Agency Health Advisory of 350 µg Mn/L.

**Results:** Estimated mean daily Mn intake from breast milk ranged from 1.2 µg Mn/kg/day (3 weeks) to 0.16 µg Mn/kg/day (18 months), with the highest intakes at the youngest age stage we considered, 3 weeks. Estimated daily Mn intake from formula products reconstituted with 0 µg Mn/L water ranged from 130 µg Mn/kg/day (3 weeks) to 4.8 µg Mn/kg/day (18 months) with the highest intakes at 3 weeks. Formula products provided 28 to 520 times greater than the mean daily intake of Mn from breast milk for the 4 age stages that we considered. Estimated daily Mn intake from formula products reconstituted with water containing 250 µg Mn/L ranged from 12 µg Mn/kg/day to 170 µg Mn/kg/day, which exceeds the United States Environmental Protection Agency Reference Dose of 140 µg Mn/kg/day for adults.

**Conclusions:** Mn deficiency is highly unlikely with exclusive breast milk or infant formula feeding, but established tolerable daily intake levels for Mn may be surpassed by some of these products when following labeled instructions.

**Highlights:**

1. Mn deficiency is unlikely with exclusive breast milk or infant formula feeding.
2. Breast milk Mn mean intake is 1.2 µg/kg/day (3 weeks)-0.16 µg/kg/day (18 months).
3. Formula Mn intake range is 130 µg/kg/day (3 weeks)-4.8 µg/kg/day (18 months).
4. Formula products reconstituted with 250 µg Mn/L water may exceed 140 µg Mn/kg/day.
5. Formula products may surpass regulatory tolerable daily intake levels for Mn.

## Background

^1^ Until recently, the effects of excess manganese (Mn) on children were unknown since studies of the neurological effects of Mn by design did not include children. Increased use of deep well water for drinking has necessitated new research on the developmental effects of common drinking water contaminants including Mn [1-6]. Recent research on Mn in drinking water has found that children who drink water with relatively high concentrations of Mn have lower intelligence quotients (IQs) and greater incidence of learning disabilities and behavioral problems than children who drink water with lower Mn concentrations [1,2,5-15]. Several recent proposals to update Mn drinking water guidelines have recognized infants as the population that is most potentially vulnerable to harm from excess exposures, but there are currently no data available concerning neurological consequences of high or low dietary post-natal Mn exposures in infants, toddlers, and young children [16-18]. Regulations regarding Mn content in infant formulas have not been updated in accordance with recent research findings on Mn toxicity and often there are no regulations for young child nutritional beverages (follow-on/-up and toddler formulas) [19].

In this study, we estimated total daily Mn exposures/kg of body weight due to breast milk and to infant formulas and young child nutritional beverage products on the market in the United States (US) and France in order to facilitate direct comparisons of Mn intakes between these feeding options. We also compared these Mn intake exposures to sufficiency and toxicity thresholds used by regulatory agencies.

## Material and methods

### Estimating mean, minimum and maximum Mn content in breast milk

To estimate the Mn content in human milk, we searched the literature for all pertinent studies published from January 1980 through December 2017 that measured Mn concentrations in human milk. Studies reporting Mn concentration in breast milk on a non-volume basis often did not include enough information for unambiguous unit conversion to a volume basis, so only results originally reported on a volume basis were included.

Searches were performed in Pubmed and Google Scholar using the terms “human milk” and “breast milk” combined with the terms “manganese” and “trace elements”. Our search identified 33 papers that reported Mn concentrations in breast milk, of which 10 were excluded because they did not report concentrations on a volume basis. Two papers that reported Mn concentrations on a volume basis specified concentrations that were orders of magnitude beyond the ranges of the other research results were excluded as well. The remaining 21 papers are summarized in Table 1.

**Table 1.**
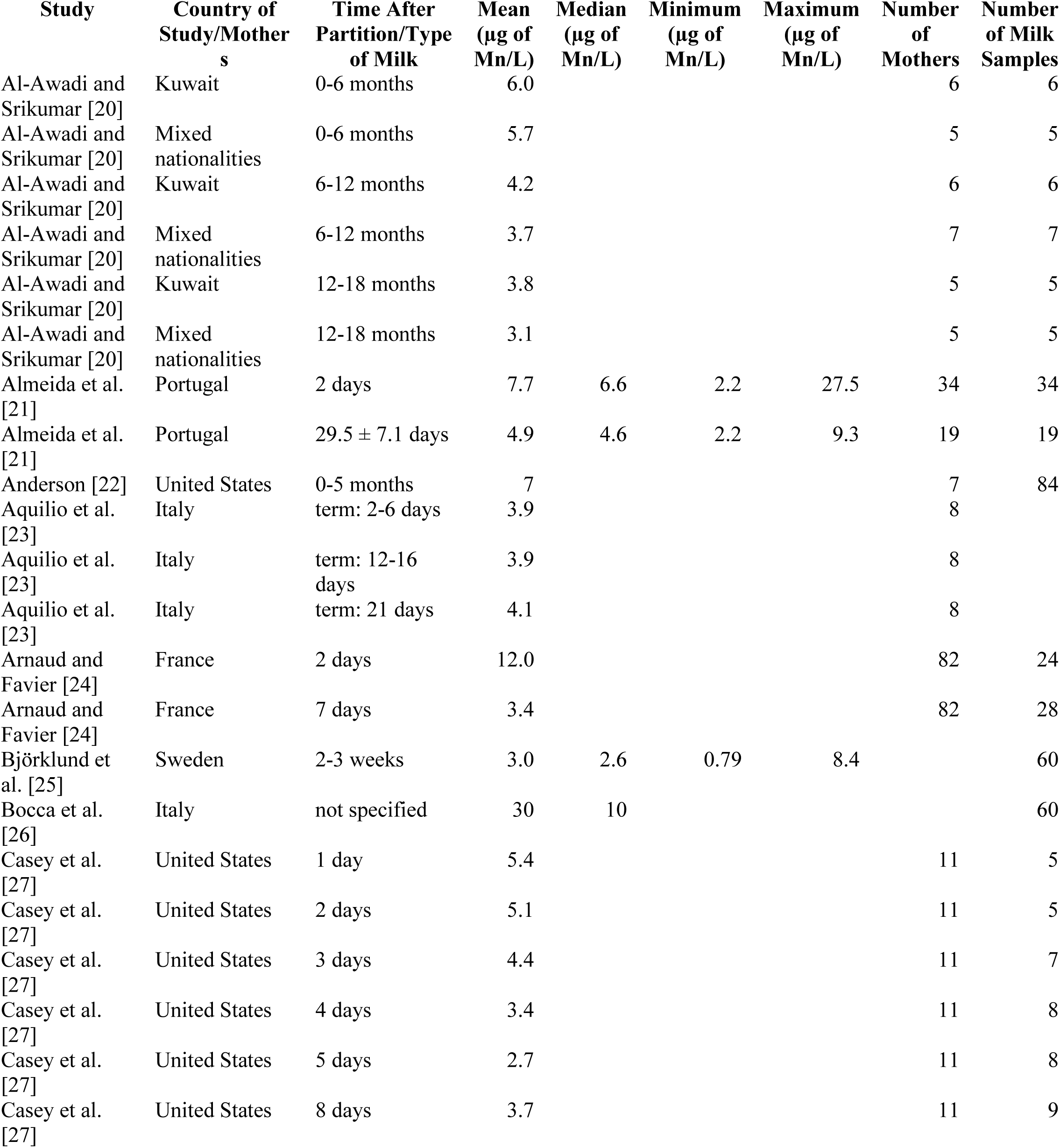
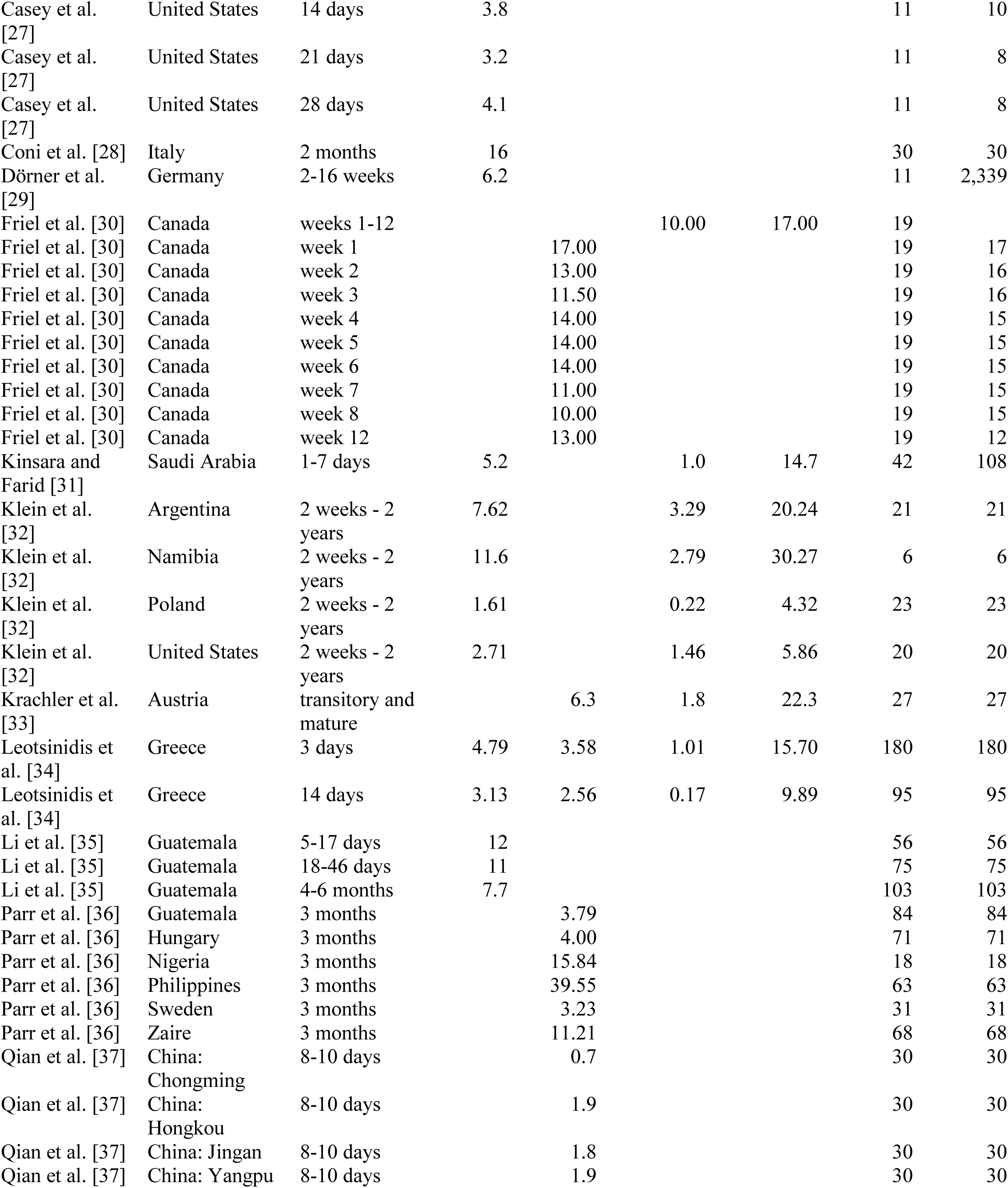
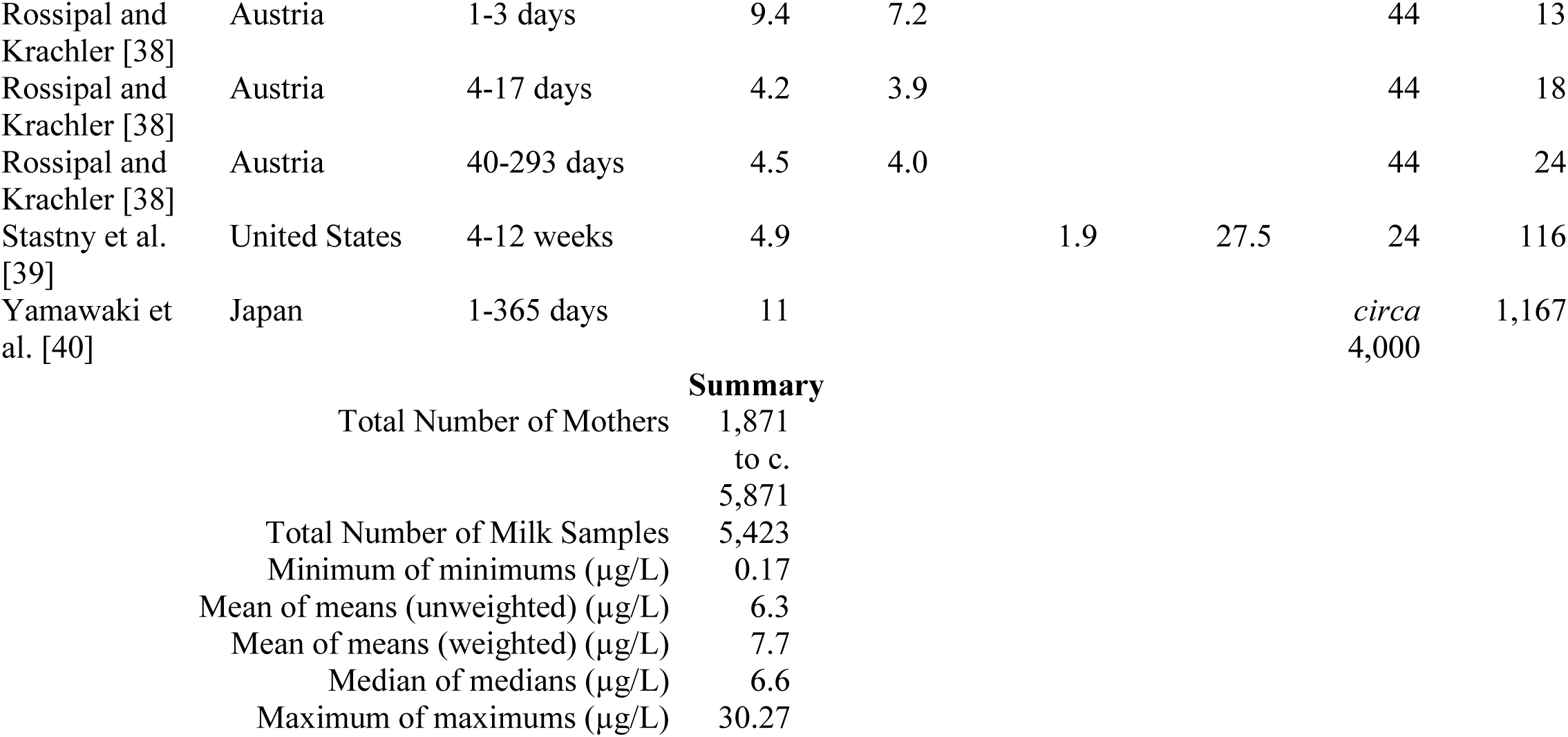
The mean, median, minimum, and maximum concentrations of manganese (Mn) in breast milk at various times after birth reported on a volume basis in the peer-reviewed literature from January 1980 through December 2017.

| Study | Country of Study/Mothers | Time After Partition/Type of Milk | Mean (µg of Mn/L) | Median (µg of Mn/L) | Minimum (µg of Mn/L) | Maximum (µg of Mn/L) | Number of Mothers | Number of Milk Samples |
| --- | --- | --- | --- | --- | --- | --- | --- | --- |
| Al-Awadi and Srikumar [20] | Kuwait | 0-6 months | 6.0 |  |  |  | 6 | 6 |
| Al-Awadi and Srikumar [20] | Mixed nationalities | 0-6 months | 5.7 |  |  |  | 5 | 5 |
| Al-Awadi and Srikumar [20] | Kuwait | 6-12 months | 4.2 |  |  |  | 6 | 6 |
| Al-Awadi and Srikumar [20] | Mixed nationalities | 6-12 months | 3.7 |  |  |  | 7 | 7 |
| Al-Awadi and Srikumar [20] | Kuwait | 12-18 months | 3.8 |  |  |  | 5 | 5 |
| Al-Awadi and Srikumar [20] | Mixed nationalities | 12-18 months | 3.1 |  |  |  | 5 | 5 |
| Almeida et al. [21] | Portugal | 2 days | 7.7 | 6.6 | 2.2 | 27.5 | 34 | 34 |
| Almeida et al. [21] | Portugal | 29.5 ± 7.1 days | 4.9 | 4.6 | 2.2 | 9.3 | 19 | 19 |
| Anderson [22] | United States | 0-5 months | 7 |  |  |  | 7 | 84 |
| Aquilio et al. [23] | Italy | term: 2-6 days | 3.9 |  |  |  | 8 |  |
| Aquilio et al. [23] | Italy | term: 12-16 days | 3.9 |  |  |  | 8 |  |
| Aquilio et al. [23] | Italy | term: 21 days | 4.1 |  |  |  | 8 |  |
| Arnaud and Favier [24] | France | 2 days | 12.0 |  |  |  | 82 | 24 |
| Arnaud and Favier [24] | France | 7 days | 3.4 |  |  |  | 82 | 28 |
| Björklund et al. [25] | Sweden | 2-3 weeks | 3.0 | 2.6 | 0.79 | 8.4 |  | 60 |
| Bocca et al. [26] | Italy | not specified | 30 | 10 |  |  |  | 60 |
| Casey et al. [27] | United States | 1 day | 5.4 |  |  |  | 11 | 5 |
| Casey et al. [27] | United States | 2 days | 5.1 |  |  |  | 11 | 5 |
| Casey et al. [27] | United States | 3 days | 4.4 |  |  |  | 11 | 7 |
| Casey et al. [27] | United States | 4 days | 3.4 |  |  |  | 11 | 8 |
| Casey et al. [27] | United States | 5 days | 2.7 |  |  |  | 11 | 8 |
| Casey et al. [27] | United States | 8 days | 3.7 |  |  |  | 11 | 9 |
| Casey et al. [27] | United States | 14 days | 3.8 |  |  |  | 11 | 10 |
| Casey et al. [27] | United States | 21 days | 3.2 |  |  |  | 11 | 8 |
| Casey et al. [27] | United States | 28 days | 4.1 |  |  |  | 11 | 8 |
| Coni et al. [28] | Italy | 2 months | 16 |  |  |  | 30 | 30 |
| Dörner et al. [29] | Germany | 2-16 weeks | 6.2 |  |  |  | 11 | 2,339 |
| Friel et al. [30] | Canada | weeks 1-12 |  |  | 10.00 | 17.00 | 19 |  |
| Friel et al. [30] | Canada | week 1 |  | 17.00 |  |  | 19 | 17 |
| Friel et al. [30] | Canada | week 2 |  | 13.00 |  |  | 19 | 16 |
| Friel et al. [30] | Canada | week 3 |  | 11.50 |  |  | 19 | 16 |
| Friel et al. [30] | Canada | week 4 |  | 14.00 |  |  | 19 | 15 |
| Friel et al. [30] | Canada | week 5 |  | 14.00 |  |  | 19 | 15 |
| Friel et al. [30] | Canada | week 6 |  | 14.00 |  |  | 19 | 15 |
| Friel et al. [30] | Canada | week 7 |  | 11.00 |  |  | 19 | 15 |
| Friel et al. [30] | Canada | week 8 |  | 10.00 |  |  | 19 | 15 |
| Friel et al. [30] | Canada | week 12 |  | 13.00 |  |  | 19 | 12 |
| Kinsara and Farid [31] | Saudi Arabia | 1-7 days | 5.2 |  | 1.0 | 14.7 | 42 | 108 |
| Klein et al. [32] | Argentina | 2 weeks - 2 years | 7.62 |  | 3.29 | 20.24 | 21 | 21 |
| Klein et al. [32] | Namibia | 2 weeks - 2 years | 11.6 |  | 2.79 | 30.27 | 6 | 6 |
| Klein et al. [32] | Poland | 2 weeks - 2 years | 1.61 |  | 0.22 | 4.32 | 23 | 23 |
| Klein et al. [32] | United States | 2 weeks - 2 years | 2.71 |  | 1.46 | 5.86 | 20 | 20 |
| Krachler et al. [33] | Austria | transitory and mature |  | 6.3 | 1.8 | 22.3 | 27 | 27 |
| Leotsinidis et al. [34] | Greece | 3 days | 4.79 | 3.58 | 1.01 | 15.70 | 180 | 180 |
| Leotsinidis et al. [34] | Greece | 14 days | 3.13 | 2.56 | 0.17 | 9.89 | 95 | 95 |
| Li et al. [35] | Guatemala | 5-17 days | 12 |  |  |  | 56 | 56 |
| Li et al. [35] | Guatemala | 18-46 days | 11 |  |  |  | 75 | 75 |
| Li et al. [35] | Guatemala | 4-6 months | 7.7 |  |  |  | 103 | 103 |
| Parr et al. [36] | Guatemala | 3 months |  | 3.79 |  |  | 84 | 84 |
| Parr et al. [36] | Hungary | 3 months |  | 4.00 |  |  | 71 | 71 |
| Parr et al. [36] | Nigeria | 3 months |  | 15.84 |  |  | 18 | 18 |
| Parr et al. [36] | Philippines | 3 months |  | 39.55 |  |  | 63 | 63 |
| Parr et al. [36] | Sweden | 3 months |  | 3.23 |  |  | 31 | 31 |
| Parr et al. [36] | Zaire | 3 months |  | 11.21 |  |  | 68 | 68 |
| Qian et al. [37] | China: Chongming | 8-10 days |  | 0.7 |  |  | 30 | 30 |
| Qian et al. [37] | China: Hongkou | 8-10 days |  | 1.9 |  |  | 30 | 30 |
| Qian et al. [37] | China: Jingan | 8-10 days |  | 1.8 |  |  | 30 | 30 |
| Qian et al. [37] | China: Yangpu | 8-10 days |  | 1.9 |  |  | 30 | 30 |
| Rossipal and Krachler [38] | Austria | 1-3 days | 9.4 | 7.2 |  |  | 44 | 13 |
| Rossipal and Krachler [38] | Austria | 4-17 days | 4.2 | 3.9 |  |  | 44 | 18 |
| Rossipal and Krachler [38] | Austria | 40-293 days | 4.5 | 4.0 |  |  | 44 | 24 |
| Stastny et al. [39] | United States | 4-12 weeks | 4.9 |  | 1.9 | 27.5 | 24 | 116 |
| Yamawaki et al. [40] | Japan | 1-365 days | 11 |  |  |  | *circa*<br>4,000 | 1,167 |

Summary
|  |  |
| --- | --- |
| Total Number of Mothers | 1,871 |
|  | to c. |
|  | 5,871 |
| Total Number of Milk Samples | 5,423 |
| Minimum of minimums ($\mu\text{g/L}$) | 0.17 |
| Mean of means (unweighted) ($\mu\text{g/L}$) | 6.3 |
| Mean of means (weighted) ($\mu\text{g/L}$) | 7.7 |
| Median of medians ($\mu\text{g/L}$) | 6.6 |
| Maximum of maximums ($\mu\text{g/L}$) | 30.27 |

We used the total number of milk samples reported for each study to calculate a weighted mean of means for studies reporting results as means [41]; studies not explicitly reporting means were not included in the weighted mean of means. Similarly, we selected the minimum of minimums and the maximum of maximums from studies reporting these values; studies not explicitly reporting minimums or maximums were not included in the minimum of minimums or maximum of maximums.

### Estimating range of Mn content in infant formulas and young child nutritional beverages available in the US and France

For the range of Mn content in infant formulas and young child nutritional beverages available in the US and France, we relied on our recent market basket survey of 44 products [42]. In this survey, we purchased 25 products in the US and 19 products in France using maximum variation sampling [43], purposively selecting products to explore the range of Mn concentrations, the potential effects of specific ingredients such as soy on Mn content, and to cover a wide range of ages. We measured the Mn content of these samples on a mass basis using particle induced X-ray emission (PIXE), Rutherford backscattering spectrometry (RBS). We then calculated µg Mn/L from the measured µg Mn/g product and the measured mass of dried product that was used to make a measured final volume of prepared product as directed by the manufacturer. Similarly, we calculated µg Mn/100 kcal using the measured µg Mn/g dried prepared product and the reported kcal/mass of dried product or final volume of prepared product from the manufacturer’s label. Since the sampling method was non-probabilistic, market extreme values for Mn concentrations (minimum and maximum) were used for comparison, but not mean values for Mn concentrations from this study [42,43].

### Selecting ages for comparison

For our estimates of Mn intake due to breast milk, infant formulas, and young child nutritional beverages, we selected 4 ages for comparison: 3 weeks, 4.25 months, 7 months, and 18 months. We selected 3 weeks since this is approximately the age when breast milk Mn concentrations stabilize after declining from initially relatively high levels (Table 1) [21,25,27,35,44,45]. We selected 4.25 months since this is the oldest age at which an infant is likely to be exclusively fed with breast milk or infant formulas [46,47] and is within the recommended ages for products labeled age stage 1 (0-6 months) from the French market or “infant formula” from the US market (0-12 months). We selected 7 months since this age is within the recommended age range of products labeled age stage 2 (6-12 months) on the French market; it is also within the age range (0-12 months) of products labeled “infant formula” from the US market. Similarly, we selected age 18 months because this age is within the recommended ages of products labeled age stages 3 or 4 (1-3 years) from the French market and products labeled “toddler formula” or “toddler powder” from the US market.

### Estimating weights of infants and young children in the US and France

Formula-fed infants have both greater intakes than breast-fed infants at all ages and greater weights at all but the very youngest ages [48-50]. For our comparisons of infants ages 3 weeks and 4.25 months, we chose to use mean body weights reported by Sievers et al. [49] since these weights were directly paired with breast milk/infant formula intake data. For ages 7 months and 18 months, we used mean body weights reported by Dewey et al. [48], since they distinguish between children who were originally primarily breast-fed or formula-fed. Dewey et al. [48] report separate mean body weights for boys and girls; we averaged these mean weights for ages 7 and 18 months. For this study, we assumed that infant and young child body weights would be similar in the United States and France because both are high income countries. Observational weight data from the 2 regions are difficult to obtain since infant weight and growth surveys are typically normative [51,52] based on non-representative populations [53], or present differences between World Health Organization (WHO) reference curves rather than mean weights of representative populations [54,55].

### Estimating daily volumes of breast milk, infant formula, or young child nutritional beverages consumed in the US and France

For estimates of intakes of breast milk or formula for ages 3 weeks and 4.25 months, we chose to use average intakes reported by Sievers et al. [49]. We converted the intakes in fluid weights [49] to fluid volumes by assuming a density of 1,031 g/L for breast milk [56], or by the average measured density of the products that we tested (mean = 1,010 g/L for US “infant formulas” and French “stage 1” products) [42].

We were not able to locate studies that reported mean breast milk or formula intakes for 7- or 18-month-olds in high-income countries by weights or volumes, so we estimated these intakes based on energy intakes for these age groups. Fantino and Gourmet [57] report % total energy intakes for infant formulas and young child nutritional beverages in the diets of formula-fed infants, 58.6% for 7-month-olds and 21.3% for 13- to 18-month-olds in France. Since separate % energy intake data were not available for infants receiving breast milk at these ages, we assumed that % energy intake of the diet due to breast milk would be equal to that of infants receiving formula.

From Fantino and Gourmet’s [57] data, a mean intake of 90._8…_ kcal/kg/day (3,004 kilojoules [kJ]/day, 7.9 kg; 4.184 kJ/kcal) [58] can be calculated for 7-month-olds. (Nonsignificant digits, such as 8…, are shown as a subscript followed by an ellipsis. These nonsignificant digits are included in all steps of a calculation to prevent rounding error.) Similarly, a mean intake of 87.4_4…_ kcal/kg/day (3,878 kJ/day, 10.6 kg) can be calculated for 13-18-month-olds [57]. We used these total kcal/kg/day intakes with the body weights reported by Dewey et al. [48] to estimate total kcal intakes/day for 7-month-olds (7.3_3…_×10^2^ kcal/day receiving breast milk, 7.6_5…_×10^2^ kcal/day receiving formula) and 18-month-olds (952._1…_ kcal/day receiving breast milk, 978._3…_ kcal/day receiving formula). We applied the % intake data devoted to infant milks to these figures, yielding 4.2_7…_×10^2^ kcal/day to breast milk and 4.5_1…_×10^2^ kcal/day to infant formula for 7-month-olds, and 202._8…_ kcal/day to breast milk and 208._3…_ kcal/day to infant formula for 18-month-olds. We used data on kcal/L for breast milk (3,103.7 kJ/L for 2-6-month-olds, 3,683.2 kJ/L for 12-39-month-olds) during prolonged lactation from Mandel et al. [59] with these energy intake figures to estimate breast milk intakes of 0.57_9…_ L/day for 7-month-olds and 0.230_3…_ L/day for 18-month-olds. For formula-fed infants, we used average energy content reported on the labels for the products we tested in our market basket survey (669._0…_ kcal/L for 7-month-olds, 621._6…_ kcal/L for 18-month-olds) together with the energy estimates due to formula, yielding estimated formula intakes of 0.67_1…_ L/day for 7-month-olds and 0.335_1…_ L/day for 18-month-olds [42].

## Results and discussion

### Manganese concentrations in breast milk

Results of our literature search for Mn concentrations in breast milk are shown in Table 1. Longitudinal studies have found that the Mn concentration of breast milk is generally highest in colostrum, declines over the first few days or weeks, then remains relatively stable in mature milk (Table 1) [21,24,25,27,35,39,40,44,45,60].

The minimum of minimums of reported Mn concentrations in breast milk is 0.17 µg Mn/L and the maximum of maximums is 30.27 µg Mn/L (Table 1). The regulatory units for infant formulas are usually given in µg Mn/100 kcals prepared formula [61-64]. The 30.27 µg Mn/L maximum of maximums concentration of Mn in breast milk expressed in IF regulatory units would be equivalent to 4.34 µg Mn/100 kcal (Equation 1). This calculation assumes an average gross energy of 697 kcal/L of breast milk at 3 months of lactation [65].

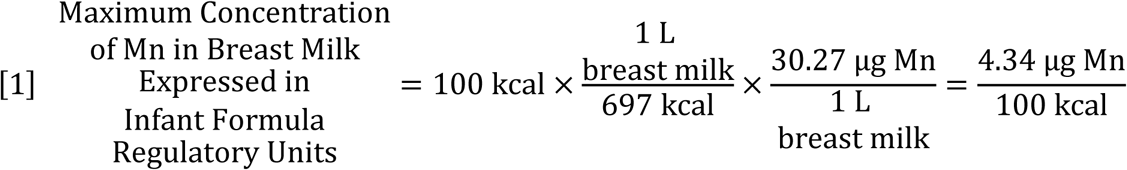

The 7.7 µg/L mean (weighted mean of means; Table 1) concentration of Mn in breast milk expressed in infant formula regulatory units would be equivalent to 1.1 µg Mn/100 kcal (Equation 2).

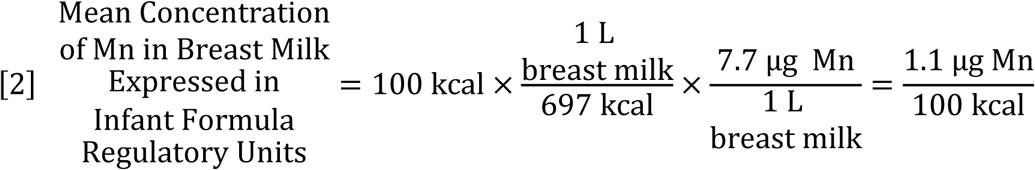

The 1 µg Mn/100 kcal minimum concentration for Mn in prepared infant formula set by the EU and the Republic of France is by design comparable to this 1.1 µg Mn/100 kcal mean concentration of Mn in breast milk (Equation 2) [63,64,66]. In contrast, the US regulatory Minimum Level of 5 µg of Mn/100 kcal Mn exceeds the 4.34 µg Mn/100 kcal maximum concentration reported in breast milk (Equation 1) [61].

### Daily Mn intake estimates from breast milk, infant formulas and young child nutritional beverages

Daily breast milk, infant formula, and young child nutritional beverage intake estimates were calculated for infants and young children of average weights from developed countries at 3 weeks, 4.25 months, 7 months, and 18 months as summarized in Table 2. These intake estimates relied on the weight, intake, and Mn data described in the Materials and Methods section (Table 1) [42,49,50,58]. These intake estimates are compared to reference weights and intake assumptions used by advisory and regulatory agencies in Table 3.

**Table 2.**
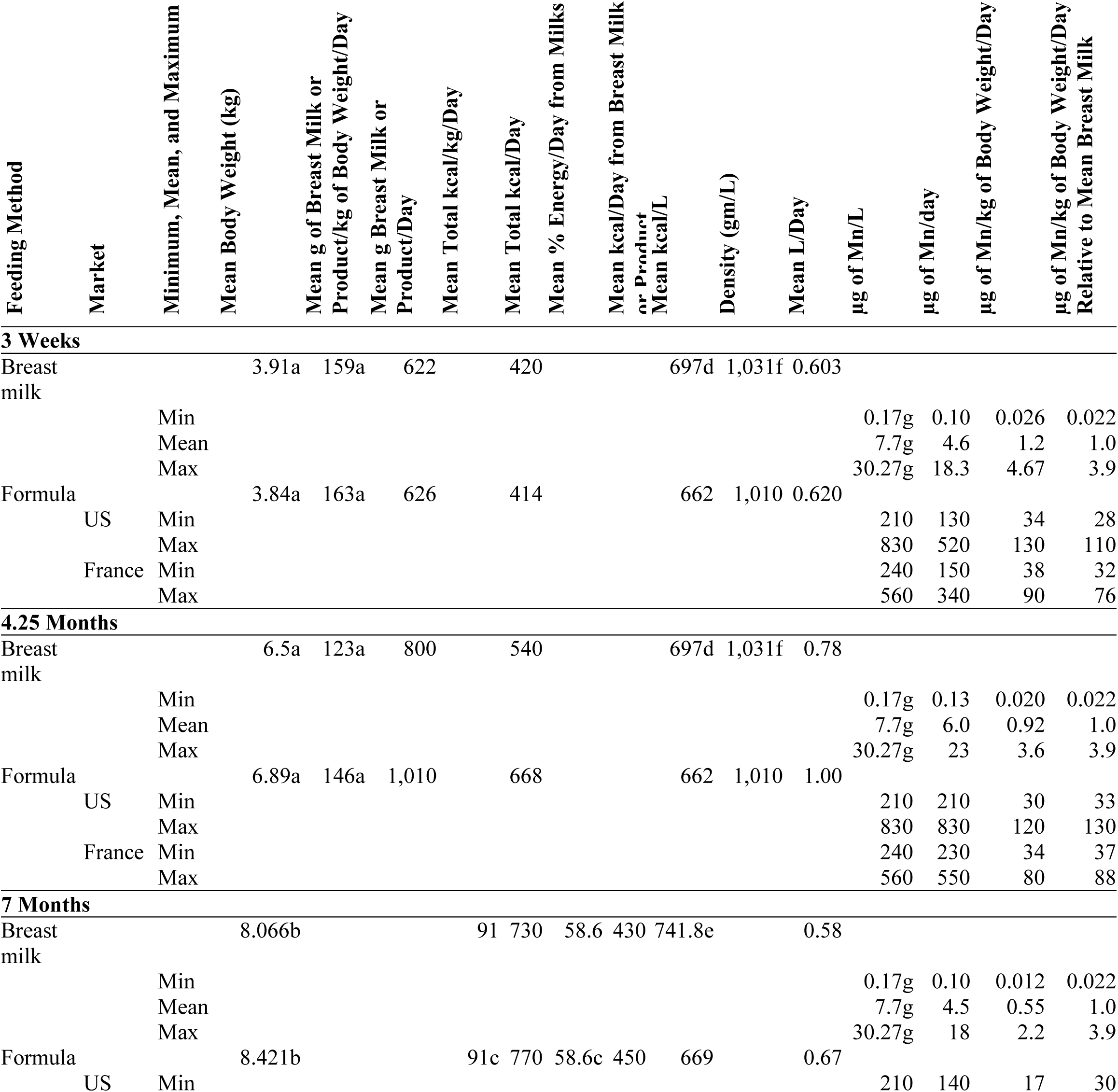
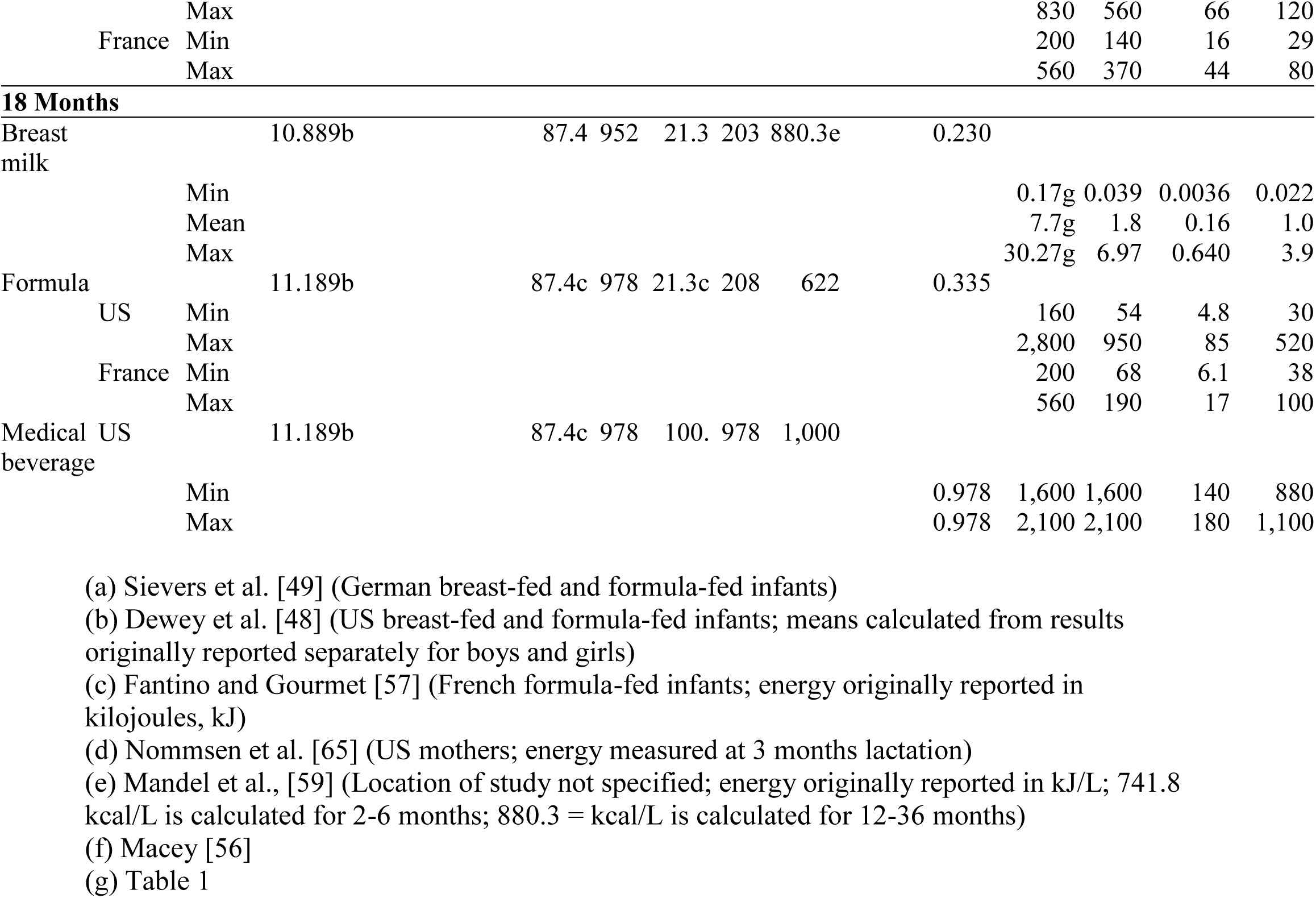
Estimated daily manganese (Mn) intakes for infants and young children at 3 weeks, 4.25 months, 7 months, and 18 months from breast milk, and minimum and maximum intakes from infant formulas and young child nutritional beverage products sampled in a market basket survey of products available on the US and French markets [42]. These estimates assume that the products are reconstituted with water containing 0 µg of Mn/L.

**Table 3.** Body weights (kg), daily breast milk or formula intakes (L/day), and daily caloric intake estimates (kcal/day) at various ages used by regulatory or advisory agencies compared to estimates from Table 2.

| Organization/Estimate from Table 2 | Age | Breast-Fed |  | Formula-Fed |  |  | Feeding Method Not Specified Body Weight (kg) |
| --- | --- | --- | --- | --- | --- | --- | --- |
|  |  | Body Weight (kg) | L/Day of Breast Milk | Body Weight (kg) | L/Day of Formula | kcal/Day |  |
| Estimates from Table 2 | 3 weeks | 3.91 | 0.603 | 3.84 | 0.620 | 414 |  |
| Environmental Protection Agency [67] | 0-<1 month | 3.40 | 0.510 |  |  |  | 4.8 |
| Environmental Protection Agency [67] | 1-<3 months | 4.93 | 0.690 |  |  |  | 5.6 |
| Estimates from Table 2 | 4.25 months | 6.5 | 0.78 | 6.89 | 1.00 | 668 |  |
| *Institut national de santé publique du Québec* (National Institute of Public Health of Quebec) [68] | <6 months |  |  |  | 1.22 (95<sup>th</sup> percentile) |  | 6.7 |
| Food and Agriculture Organization/World Health Organization (Codex Alimentarius) [69] | 0-6 months |  |  | 5 | 0.750 | 500 |  |
| European Food Safety Authority [70] | 0-6 months |  | 0.8 |  |  | 500 |  |
| *Agence Française de Sécurité Sanitaire des Aliments* (French Food Safety Agency) [71] | 0-6 months |  |  |  | 0.420 |  |  |
| Institute of Medicine [72] | 0-6 months |  | 0.78 |  |  |  |  |
| Environmental Protection Agency [67] | 3-<6 months | 7.00 | 0.770 |  |  |  | 7.4 |
| Institute of Medicine [72] | 6 months |  |  |  |  |  | 7 |
| Estimates from Table 2 | 7 months | 8.066 | 0.58 | 8.421 | 0.67 |  |  |
| Environmental Protection Agency [67] | 6-<12 months | 7.5 | 0.620 |  |  |  | 9.2 |
| Institute of Medicine [72] | 7-12 months |  | 0.6 |  |  |  |  |
| Institute of Medicine [72] | 12 months |  |  |  |  |  | 9 |
| Environmental Protection Agency [67] | 1-< years |  |  |  |  |  | 11.4 |
| Estimates from Table 2 | 18 months | 10.889 | 0.230 | 11.189 | 0.335 |  |  |
| Institute of Medicine [72] | 1-3 years |  |  |  |  |  | 13 |

Although we developed our breast milk intake estimates for ages 7 and 18 months based on weight and energy intake data rather than direct measurements since breast milk intake data were not readily available for infants and children in the developed world at these ages, our estimates appear to be within range or close to the ranges reported for similarly aged groups in the literature. Our breast milk intake estimate for 7-month-olds (0.58 L/day) is within the observed range (321-641g/day = 0.318-0.634 L/day) for 6-9 month-olds reported by a 1985 international survey of breast feeding by the WHO [73] but higher than the mean for 24-hour recall reports for 6-11.9-month-olds from a 2016 US feeding survey (564.7 g/day = 0.5591 L/day) [74]. For 18-month-olds, our breast milk intake estimate (0.23 L/day) is slightly lower than the minimum (294 g/day = 0.291 L/day) reported for 18-month-olds in developing countries [73] and also lower than the mean for recent 24-hour recall reports for 12-23.9-month-olds in the US (322.4 g/day = 0.3192 L/day) [74]. Thus, our breast milk intake for 18-month-olds based on weight and energy intakes may underestimate actual breast milk intake for this age group by 21%-28%.

We combined our estimates for weight and daily breast milk or formula intake with the minimum, mean, and maximum concentrations of Mn in breast milk (Table 1), and the products measured in our market basket survey to estimate intakes of Mn/kg/day due to breast milk, infant formula, or young child nutritional beverage products (Equations 1 and 2, Table 2, and Figure 1) [42]. Estimated daily intake of Mn due to breast milk/kg body weight was highest at the youngest age that we examined (3 weeks) and declined as age and total body weight increased (Mn/kg/day at 18 months). For infant formulas and young child nutritional beverage products, estimated daily intake of Mn due to these products was similarly highest at the 3-week-stage. Minimum intake of Mn due to these products was at least 28 times that of the mean Mn intake due to breast milk at each corresponding age (Table 2 and Figure 1).

**Fig. 1.**
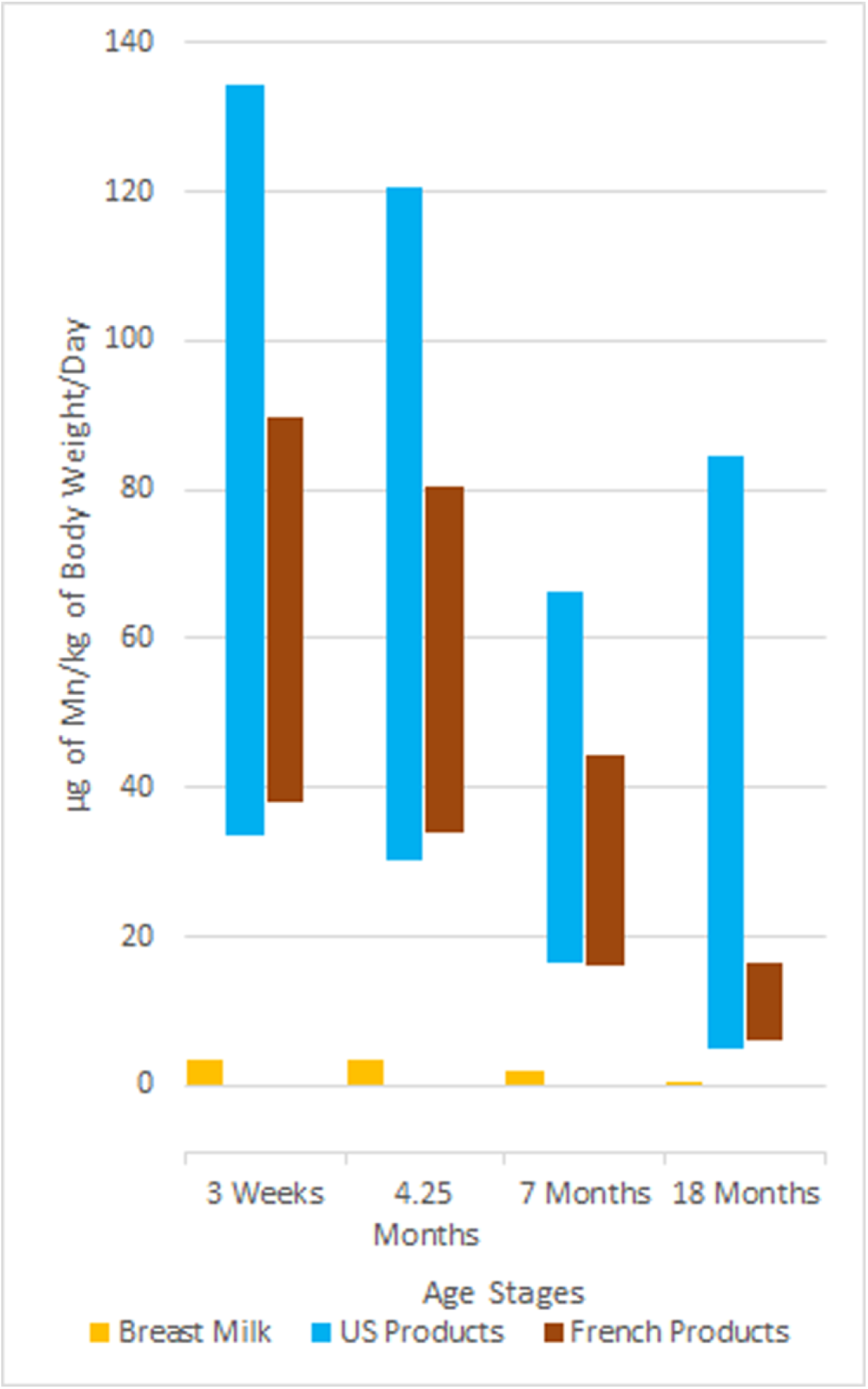
Estimated ranges of intakes of Mn/kg of body weight/day at 3 weeks, 4.25 months, 7 months, and 18 months for breast milk, United States (US) products, and French products.

Although it has been noted that “human milk composition shows remarkable variation” [66], the range of Mn concentration in breast milk is both lower and narrower than that of infant formulas [75]. The intake of Mn per day from breast milk with the maximum reported concentration of Mn is not more than 4 times the daily intake from breast milk with the mean reported concentration of Mn (Tables 1 and 2). In contrast, for infant formulas and young child nutritional beverage products, the daily intake of Mn/kg body weight from the products that we surveyed was between 28 and 520 times the daily intake of Mn/kg body weight of breast milk with the mean reported concentration of Mn (Tables 1 and 2) [42].

### Effect of water Mn content on Mn exposures due to infant formulas and young child nutritional beverages

Infant and toddler formula powders are often reconstituted with tap water and sometimes with mineral water [76,77]. While most tap water in the US and France has a Mn concentration below 50 µg/L, a level referenced by some drinking water guidelines due to aesthetic concerns such as staining laundry or plumbing fixtures [78,79], water from some sources, especially groundwater, may contain considerably higher concentrations of Mn [79-81]. Water from public sources occasionally exceeds the United States Environmental Protection Agency (U.S. EPA) Lifetime Health Advisory (HA) for Mn in drinking water of 300 µg/L [79,83]. Mineral water may also contain Mn in excess of 400 µg/L [76].

To estimate the daily Mn intake for formula-fed infants and young children consuming products made not with ultra-trace elemental analysis grade water with 0 µg/L Mn (as is typically used when performing elemental analysis of infant formulas), but rather with tap or mineral water containing appreciable Mn, we considered the hypothetical case of water containing 250 µg/L Mn, which has a relatively high Mn content compared to the stipulated aesthetic acceptability thresholds of 50 µg/L [78,79,85] to 100 µg/L [85], but is lower than the U.S. EPA HA of 300 µg/L [79] or the WHO health-based value (HBV) of 400 µg/L [85]. Although most consumers have access to water with less than 50 µg Mn/L, some regularly drink water with a relatively high Mn content [80]. In our hypothetical case of water with 250 µg Mn/L, the Mn content of the water is within health-based guidelines and regulations, so consumers might assume that it should be safe for all purposes, including feeding infants and young children.

Following the instructions printed on product labels for reconstituting the formula powders with water, we measured the mass of powder needed to make 1 “batch” of prepared product (g) as specified on the product labels, added the exact volume of water for 1 batch as specified on the labels (mL), and measured the final volume of prepared product (mL). Measurements were in triplicate and averaged for each of the 42 powdered products in this study; the remaining two products were already liquids (Table 4). We then calculated the mass of Mn contained in the product powder used to make 1 batch and the mass of Mn in an aliquot of water used to make 1 batch (µg), assuming that the water had an Mn concentration of 250 µg/L, and summed the masses of Mn from these 2 sources to calculate the total mass of Mn contained in a batch of product prepared with water with 250 µg Mn/L Mn (Table 4). Finally, we converted the various batch volumes to liters and applied the body weight (kg) and consumption (L/day) estimates for ages 3 weeks, 4.25 months, 7 months, and 18 months from Table 2. We found that the estimated total intakes of Mn may be more than double compared to when the products are reconstituted with 0 µg of Mn/L water (Tables 2 and 4).

**Table 4.**
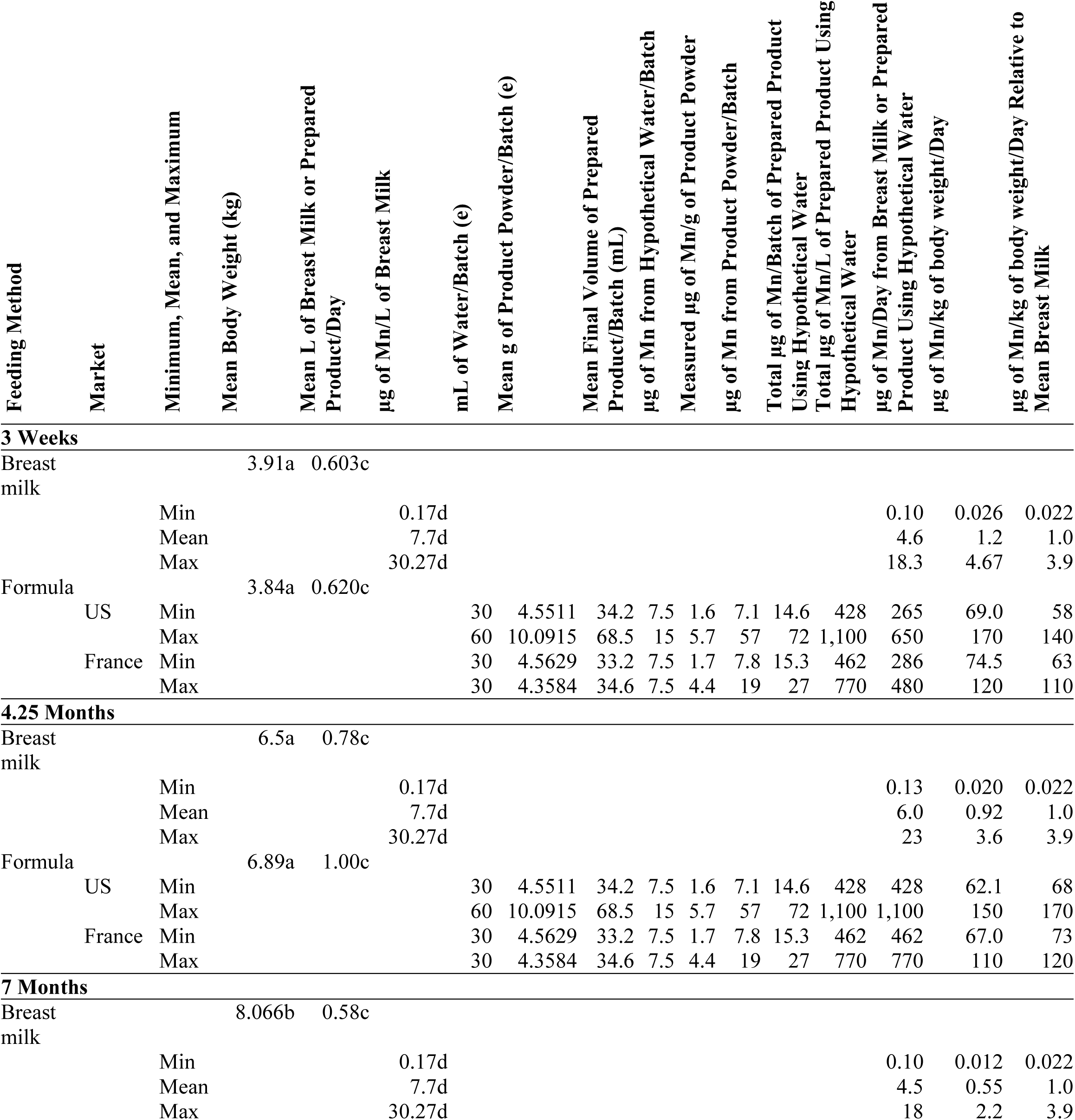
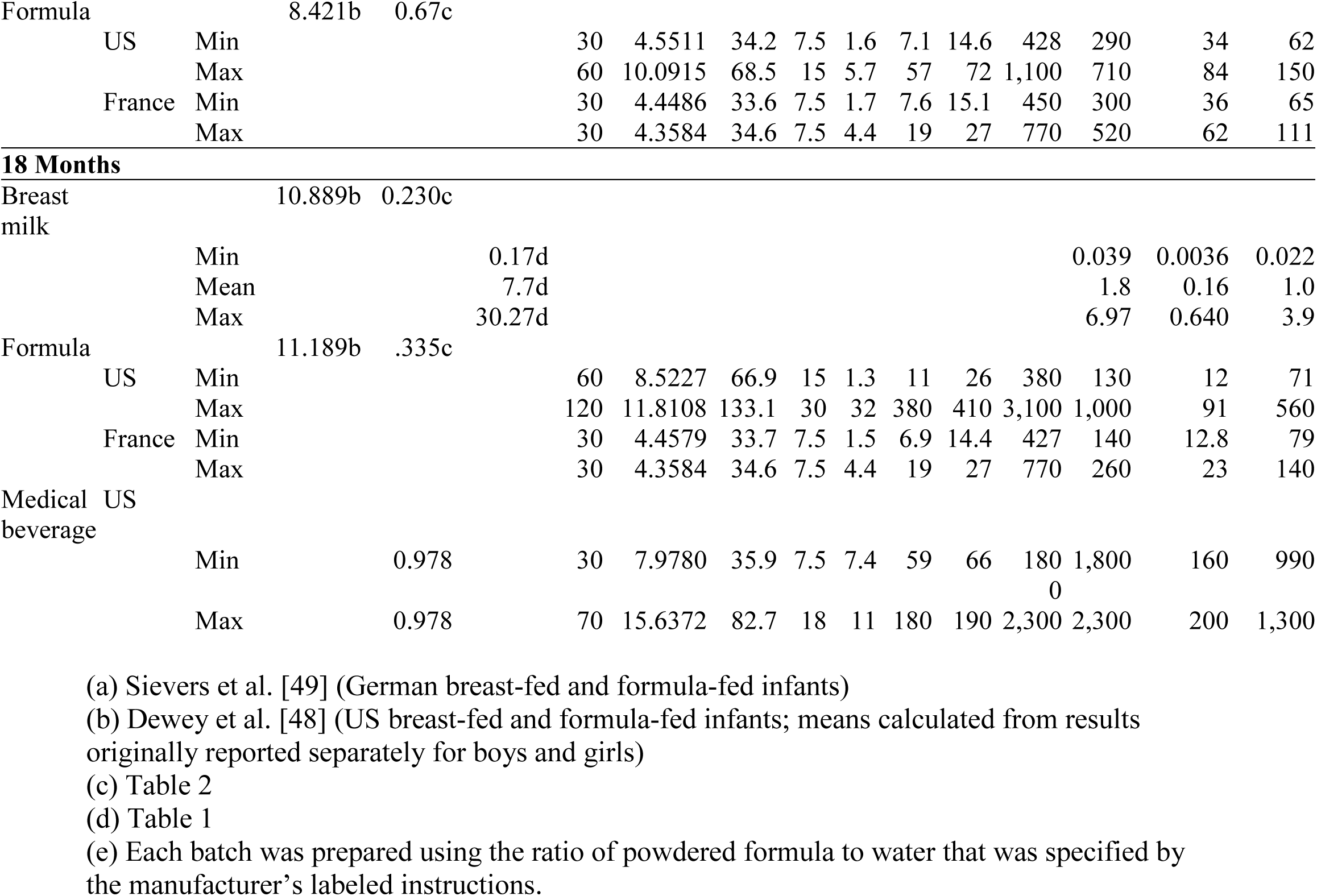
Estimated daily manganese (Mn) intakes for infants and young children at 3 weeks, 4.25 months, 7 months, and 18 months from infant formulas and young child nutritional beverage product powders reconstituted with hypothetical water containing 250 µg of Mn/L.

The U.S. EPA HA for Mn in drinking water of 300 µg/L is based on a Reference Dose (RfD) of 140 µg Mn/kg/day [79], a dose which is surpassed by the infant formulas on the US market with the highest Mn contents when reconstituted with high Mn containing waters (e.g. 250 µg/L) for 3-week-old and 4.25-month-old infants (Table 4). Notably, 2 US products labelled for use by infants ages 0 and older would exceed this RfD of 140 µg Mn/kg/day if prepared with water containing 250 µg Mn/L (Table 4) [42]. Of these 2 products, one was an infant medicinal product intended for complete nutrition and the other was a soy-based infant formula.

It must also be kept in mind that the U.S. EPA RfD of 140 µg Mn/kg/day was based on dietary studies in adults with an assumed mean body weight of 70 kg [79, 86-88] and may not be sufficiently low to protect infants whose Mn excretion capacity is not fully mature [10,11,18,89].

In sum, when infant formula powders are reconstituted with water with a Mn concentration lower than the WHO HBV of 400 µg/L [85,90-93] or the U.S. EPA HA of 300 µg/L [79], the resulting prepared formulas may lead to total daily intakes of Mn in excess of established regulatory tolerable intake levels (Table 4). Moreover, the established tolerable intake levels that we have used for comparison are based on adult physiology and do not take into account the special vulnerabilities of young infants. Tolerable intake levels for infants would most likely be set lower than those that might be extrapolated by weight from those set for adults [18].

### Essentiality of Mn, absorption, and excretion

Manganese is required for numerous essential enzymes, including the antioxidant metalloenzyme manganese superoxide dismutase (MnSOD) [94]. Manganese is provided in ordinary diets by leafy greens, nuts, grains, and animal products [90,91]. It is possible that Mn may be more absorbable from water than food, although this question has not been studied directly [2,95]. Although some research originally suggested that Mn is more absorbable from human breast milk than from cow’s milk [96], the present consensus is that there do not appear to be large differences in bioavailability of Mn in infants between breast milk and infant formula [66,97]. Fractional absorption of Mn in adults is usually assumed to be about 1-5% [97,98]; these figures are based on a rat study [99] but are comparable to those reported for fractional absorption from Mn supplements in humans [100].

In adults, Mn is excreted by the biliary system; however, the biliary excretion mechanism is not yet mature in neonates, leading to higher Mn retention at the youngest ages [10,11,89]. In experimental animal studies, neonatal rats fed supplemental Mn showed increased brain dopamine levels and changes in behavior consistent with neurodevelopmental damage [101,102]. These facts suggest the potential for excess manganese exposure and consequent neurodevelopmental damage in infants.

### Manganese deficiency is highly unlikely with either exclusive or partial formula-feeding

Symptoms of overt Mn deficiency in animals or humans have only been reported when diets are severely constrained, such as with confined farm or laboratory animals fed controlled rations low in Mn content. Laboratory and farm animals fed Mn-deficient diets exhibit structural abnormalities, adverse reproductive and fertility effects, glucose intolerance, ataxia, and hearing or vision deficits [103-107]. In humans, Mn deficiency has never been noted among people consuming ordinary diets; it has only manifested after extended periods of controlled experimental diets with very low Mn content or years of total parenteral nutrition (TPN) without Mn supplementation [94]. We have not found any studies reporting that supplementation with specifically Mn without simultaneous supplementation of other minerals to be beneficial for humans consuming ordinary diets.

Infants who consume foods in addition to formula (complementary foods) ingest sufficient Mn through the complementary foods in their diet [72]. Infants generally begin consuming complementary foods between ages 4-6 months [77], so Mn deficiency is highly unlikely after 6 months of age in either breast-fed or formula-fed children. Mn deficiency has never been reported in exclusively breast-fed infants, so it is assumed that breast milk provides sufficient Mn for development in young infants, 0-6 months [72].

The US Institute of Medicine (IOM) has set an Adequate Intake (AI) for 0-6 month-old-infants of 3 µg Mn/day based on an estimated Mn concentration in breast milk of 3.5 µg/L (reference weights not specified) [72]. All products from our market basket survey supplied more than the AI for Mn; the minimum concentration of Mn in the infant products that we tested would supply 130 µg Mn/day for 3-week-olds when prepared with water that contained no Mn, approximately 43 times the AI suggested by the IOM for 0-6-month-olds (Table 2) [42,72].

For older infants (7-12 months), the IOM assumes that the infants will include ordinary foods in their diet and extrapolates the adult AI to these older infants according to weight (7 kg) for an AI of 600 µg Mn/day (Table 3) [72]. According to the estimated intakes in Table 2, breast milk with a mean Mn concentration of 7.7 µg/L would supply approximately 0.74% of this AI with an intake of 0.58 L of breast milk/day, with the remainder of the AI of Mn to be supplied by complementary foods. Since Mn deficiency has never been reported for 7-month-old breast-fed infants, the daily intake of Mn through breast milk and complementary foods is likely sufficient. In contrast, the infant formula product that we tested with the minimum Mn concentration for this age stage would supply approximately 23% of the AI with an intake of 0.67 L of formula/day. The infant formula product with the minimum Mn concentration for this age group in our market basket survey supplies 29 times as much Mn as breast milk, so the total intake of this product together with complementary foods will likely be more than sufficient (Table 2) [42].

The IOM sets the AI for 1-3 years at 1,220 µg Mn/day based on median intake of dietary studies [72]. According to the estimated intakes in Table 2, breast milk with a mean concentration of 7.7 µg Mn/L would supply 0.15% of this AI with an intake of 0.230 L breast milk/day for 18-month-olds. In contrast, the product from our market basket survey with the minimum Mn concentration for this age stage would supply 4.4% of this AI with an intake of 0.335 L formula/day for 18-month-olds (Table 2) [42]. Since Mn deficiency has never been reported for 18-month-old breast-fed infants, the daily intake of Mn through breast milk and complementary foods is likely sufficient. The product from our market basket survey with the minimum Mn concentration for this age group supplies 30 times as much Mn as breast milk, so the total intake of this product together with complementary foods will likely be more than sufficient (Table 2) [42].

Thus, all of the products from our market basket survey would supply much more Mn per day and a greater proportion of the daily AI than breast milk at all of the age stages that we examined [42]. Most products that we tested were supplemented with Mn, but the unsupplemented dairy infant formula with the minimum Mn concentration (230 µg/L) would supply 29 times the daily Mn intake as breast milk for a 3-week-old and approximately 47 times the AI for Mn for this age stage. Cow and goat milks generally contain higher concentrations of Mn than human breast milk (Table 1) [20,108,109], and soy and rice proteins have even higher Mn concentrations [110,111], so it is highly unlikely that there is ever a need for Mn supplementation of infant formulas or young child nutritional beverage products in order to prevent Mn deficiency or achieve nutritional adequacy.

### The toxicity of manganese by ingestion exposure

The dietary studies quantifying Mn exposure used by regulatory agencies to set the Mn ingestion guidelines that are currently in force were simply estimates of intakes without any measurements of health outcomes [72,79,90,112-114]. Recent research has suggested that high dietary intakes of Mn are associated with increased inflammatory markers in elderly men [115]. Chronic consumption of dietary supplements containing excessive levels of Mn has led to the development of Parkinson’s Disease, CNS disturbances, and dementia in adults [116-118]. Possible health effects of ingestion of high levels of Mn have not been studied in infants, toddlers, or young children. However, infants who receive parental nutrition (PN) for extended periods are prone to excess accumulation of Mn in the basal ganglia related to their intravenous Mn intake, which can be seen with shortened T1 relaxation (T1R) times using magnetic resonance (MR) relaxometry [119]. It is not known whether dietary exposure alone could cause effects such as those found with pediatric PN patients, but the nature of the neurological effects observed with Mn exposures through drinking water in school-aged children, as well as the demonstrated neurological effects of ingestion exposures in adults suggest the potential for neurological effects due to excess dietary exposures in infants, toddlers, and young children [120,121]. Since biomarkers that reliably correlate with Mn exposures have not yet been identified [119,122-124], regulatory toxicity thresholds for Mn by ingestion are generally based on exposures rather than biomarkers [72,79,85].

### Infant formulas and young child nutritional beverage products must be compared to tolerable intake levels, NOT drinking water guidelines

Organizations which disseminate food standards or guidelines have established tolerable intake levels for certain common nutrients or contaminants in foods, such as arsenic in rice [125]. However, such standards or guidelines are not available for many of the nutrients or contaminants that commonly occur in food. In contrast, standards or guidelines may be available for these contaminants in drinking water. In order to determine whether beverage products exceed published tolerable intake levels, some researchers have compared their measured concentrations of nutrients or contaminants directly to drinking water standards or guidelines since the units are conveniently expressed in mass of nutrient or contaminant per volume of drinking water. Numerous examples of this practice can be found in the literature by searching for individual contaminants (eg. arsenic or lead) and specific beverage types (eg. wine or apple juice). However, this practice is highly problematical and incorrect since health-based drinking water standards or guidelines always incorporate intake assumptions specific to drinking water that are inappropriate for dietary exposures.

For example, when deriving a drinking water guideline to protect against adverse health effects in adults, the WHO usually assumes an intake of 2 L drinking water per day by a 60 kg adult and typically a 20% allocation factor for drinking water [92]. That is, 20% of the exposure is typically estimated to come from drinking water and the remaining 80% is estimated to come from all other sources, such as ingesting food or soil.

Using drinking water standards or guidelines to directly determine the potential toxicity of a chemical in a nutrient-containing liquid, as has been frequently done in prior beverage studies, is invalid since the intake from a nutrient-containing liquid is dietary and must be accounted for in the dietary portion of the exposure allocation factor, which is separate from the intake from drinking water. Also, the standard body weight and intake assumptions used in the derivation of a drinking water standard or guideline may yield a ratio that is inappropriate for liquids that are not drinking water, especially for nutrient-containing liquids consumed by infants or young children.

Nevertheless, when standards or guidelines for a nutrient or contaminant in food are not available, tolerable intake levels that could be used for comparison of measured concentrations may be identified through examination of the *derivations* of drinking water standards or guidelines. This is because, in addition to the standard assumptions specific to drinking water, all health-based drinking water standard or guideline derivations refer to a tolerable intake level of the toxic substance expressed in terms of mass of substance/kg body weight/day.

Since tolerable intake levels are expressed in mass of substance/kg body weight/day, to compare measured concentrations from nutrient-containing liquids to these tolerable intake levels, one must estimate the body weight of the consumer, the total intake of the nutrient-containing liquid, and the percent of total intake of the nutrient or contaminant that this dietary intake represents. Weight and intake estimates for infant formula consumption at various ages are provided in Table 2. In the case of a dietary liquid used for complete nutrition such as infant formula, the infant formula as prepared may provide 100% of the total ingestion intake.

Keeping in mind that drinking water guidelines or standards must *never* be used for direct comparison to nutrient-containing liquids such as infant formula, a number of international and national organizations have established tolerable intake levels for Mn in drinking water whose *derivations* could be used to estimate equivalent tolerable intake levels in infant formulas. Calculating such equivalences requires the use of the body weights and intake estimates specific for the population and substance (see above: *Estimating weights of infants and young children in the US and France*, and *Estimating daily volumes of breast milk, infant formula, or young child nutritional beverages consumed in the US and France*, Table 2). In particular, the following organizations have established tolerable intakes levels for the ingestion of Mn:

- WHO: No Observed Adverse Effect Level (NOAEL) of 11 mg Mn/day for adults based on adult dietary exposures [91]
- IOM: Upper Limit (UL) of 2 mg Mn per day for children ages 1-3 based on adult dietary exposures [72]
- US EPA: Reference Dose (RfD) of 140 µg Mn/kg/day based on adult dietary exposures [79]
- *Agence nationale de sécurité sanitaire de l’alimentation, de l’environnement et du travail* (*ANSES*, French Agency for Food, Environmental and Occupational Health & Safety): *dose journalière admissible* (*DJA*, tolerable daily intake or TDI) of 55 μg Mn/kg/day for infants based on water exposures in rats [16]

### The WHO No Observed Adverse Effect Level for the ingestion of manganese in water

The United Nations Food and Agriculture Organization (FAO) and the Joint Expert Committee for Food Additives (JECFA) have not established a tolerable intake level for Mn in food. However, the WHO has published an HBV for Mn in drinking water [85] assuming a NOAEL of 11 mg Mn/day for adults based on adult dietary exposures [91]. In order to compare the Mn content of infant formulas and young child nutritional beverages to this reference value, we must factor in differences in body size and daily consumption amounts before we can estimate an equivalence for Mn content per L.

When using this NOAEL, the WHO assumes that it applies to 60 kg adults [91], which would yield an upper tolerable limit of 180 µg Mn/kg/day for dietary intake (Equation 3):

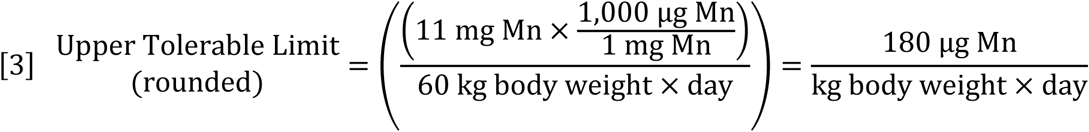

For a 3-week-old formula-fed infant weighing 3.84 kg (Table 2), 180 µg Mn/kg/day would correspond to a daily intake of 700 µg Mn per day as follows (Equations 3 and 4).

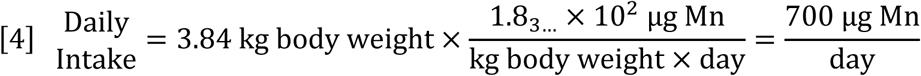

With an intake of 0.619_7…_ L of prepared product per day for a 3-week-old (Table 2), 700 µg Mn/day would be provided by an infant formula with 1,100 µg Mn/L of product as follows (Equation 5).

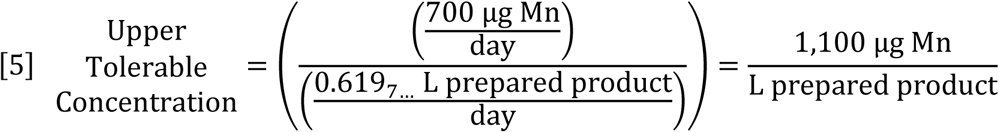

In our market basket survey of products on the US and French markets, none of the infant formulas exceeded this 1,100 µg Mn/L upper tolerable concentration for 3-week-olds or the corresponding 180 µg Mn/kg/day upper tolerable limit (Equations 3 and 5) [42,91].

### The IOM Tolerable Upper Intake Level (UL) for the ingestion of manganese

In 2001, the US IOM set a Tolerable Upper Intake Level (UL) of 11 mg Mn/day “for adults based on a no-observed-adverse-effect level for Western diets” [72]. The IOM used their 11 mg Mn/day UL for adults to develop ULs for the ingestion of Mn in food by children at different ages [72]. The US Agency for Toxic Substances and Disease Registry (ATSDR) has adopted the IOM [72] UL of 11 mg Mn/day for 70 kg adults as an interim guidance value for risk assessments of oral exposure to inorganic forms of Mn [126].

In developing its ULs for Mn, the IOM considered adults, infants, children, and adolescents separately [72]. The IOM states, “For infants, the UL was judged not determinable because of lack of data on adverse effects in this age group and concern about the infant’s ability to handle excess amounts. To prevent high levels of manganese intake, the only source of intake for infants should be from food or formula” [72].

For children 1-3 years old, the IOM assumed a 13 kg body weight and extrapolated a 2 mg Mn per day (2,000 µg Mn/day) UL to 1 significant figure as follows (Equation 6) [72].

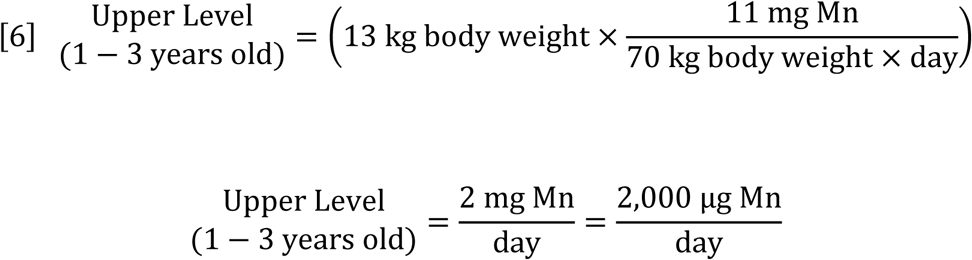

This UL applies to all dietary Mn intake, including Mn from complementary foods as well as young child nutritional beverages. Fantino & Gourmet [57] reported that formulas provide an average 21.3% of the total daily energy intake for 1-3-year-old children. If the contribution to total Mn intake from formulas is proportional to the contribution to total energy intake, then formulas should provide no more than 21.3% of the IOM 2,000 µg Mn/day UL (Table 2). This would yield a 1,200 μg Mn/L of prepared product upper tolerable concentration for 1-3-year-old children consuming complementary foods as follows (Equations 6 and 7).

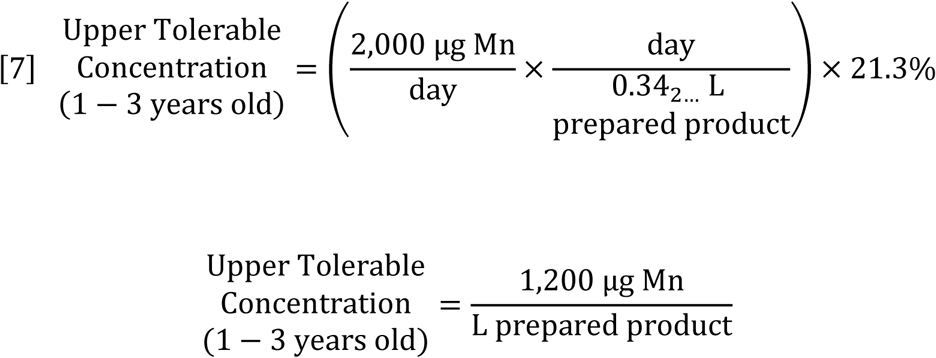

In our market basket survey of infant formulas and young child nutritional beverages on the US and French markets, none of the 19 products purchased in France exceeded this 1,200 µg Mn/L prepared product upper tolerable concentration [42]. One of the products purchased in the US, a toddler formula, contained 2,800 µg Mn/L of prepared product, exceeding this proportional upper tolerable concentration by more than 2 times [42].

Products labelled as providing complete nutrition for young children must be considered separately since their intake estimates differ substantially from ordinary infant or follow-up formulas. These products are used in cases of medical necessity and are expected to supply 100% of the child’s nutritional intake. The upper tolerable concentration for a 1-3-year-old child consuming a nutritionally complete medical beverage was estimated by using the IOM 2,000 µg Mn/day UL for children ages 1-3 years [72], the assumed 13 kg of body weight for this age group [72], and an energy requirement of 87.4_4…_ kcal/kg/day for 18-month-olds (Table 2) [57] as follows (Equation 8).

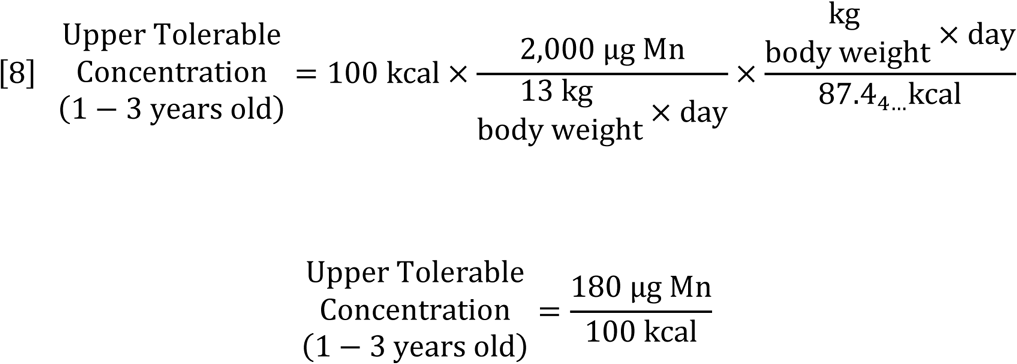

One of the 2 nutritionally complete medical beverage products labeled for ages 1 year and older that we purchased in the US for our market basket survey exceeded this upper tolerable concentration of 180 µg Mn/100 kcal; it contained 240 µg Mn/100 kcal when prepared according to manufacturer’s instructions [42].

### The U.S. EPA RfD for the ingestion of manganese

The U.S. EPA RfD of 140 µg Mn/kg/day was derived from a 10 mg Mn/day NOAEL based on dietary intakes [79]. None of the 20 products labeled for use by children under 1 year old in our market basket survey of infant formulas and young child nutritional beverages available in the US and France would result in an intake above this 140 µg Mn/kg/day RfD if prepared with water containing 0 µg Mn/L (Table 2) [42]. However, of the 2 nutritionally complete medical products labelled for use by children ages 1 year and older, one would equal the 140 µg Mn/kg/day U.S. EPA RfD and one would exceed it, providing 200 µg Mn/kg/day when used for complete nutrition by an 18-month-old and prepared with water containing 0 µg Mn/L (Table 2) [42].

If prepared with water containing 250 µg Mn/L, 2 of the products purchased in the US labelled for use by infants ages 0 and older would exceed the RfD of 140 µg Mn/kg/day when consumed by 3-week-old infants, as would both nutritionally complete medical products labelled for ages 1 year and older when used to provide all nutritional requirements for 18-month-olds (Table 4). Of the 2 products for infants exceeding the 140 µg Mn/kg/day RfD, one was an infant medicinal product intended for complete nutrition and the other was a soy-based infant formula.

### The EU has not set tolerable intake levels for the ingestion of manganese

Repeating the conclusions of the Scientific Committee on Food (SCF), the European Food Safety Authority (EFSA) states “Oral intake of manganese despite its poor absorption in the gastrointestinal tract has also been shown to cause neurotoxic effects. The limitations of the human data and the non-availability of NOAELs for critical endpoints from animal studies produce a considerable degree of uncertainty. Therefore, an upper level cannot be set” [127,128]. EFSA reiterates the SCF [128] conclusion, “oral exposure to manganese beyond the normally present in food and beverages could represent a risk of adverse health effects without evidence of any health benefit” [127].

In European Council Directive 98/83/EC “on the quality of water intended for human consumption”, a 50 μg Mn/L indicator parameter was set for Mn [129]. Indicator parameters are based on aesthetic concerns such as taste or color and are not health-based, so no health-based tolerable intake level for Mn can be determined from this parameter.

### The *Agence nationale de sécurité sanitaire de l’alimentation, de l’environnement et du travail* (ANSES) proposed maximum tolerable health value for manganese

The *République Française* (Republic of France) set national drinking water standards in 2001 [130]. Following the European Council Directive 98/83/EC, only an indicator value of 50 μg/L was set for Mn, and no health-based standards for Mn in water were adopted [129,130].

However, in 2018, the *Agence nationale de sécurité sanitaire de l’alimentation, de l’environnement et du travail* (ANSES) proposed a *valeur sanitaire maximale admissible* (maximum tolerable health value) for Mn in drinking water of 60 μg Mn/L [16]. Following the *Institut de santé publique du Québec* (INSPQ, Institute of Public Health of Quebec), ANSES assumed a lowest observed adverse effect level (LOAEL) of 25 mg/kg/day from studies involving juvenile rats ingesting Mn in water, modified by a total uncertainty factor of 450, yielding a *dose journalière admissible* (*DJA*, tolerable daily intake or TDI) of 55 μg Mn/kg/day [16,18].

For a 3-week-old formula-fed infant weighing 3.84 kg (Table 2), 55 µg Mn/kg/day would correspond to a daily intake of 210 µg Mn per day as follows (Equations 9).

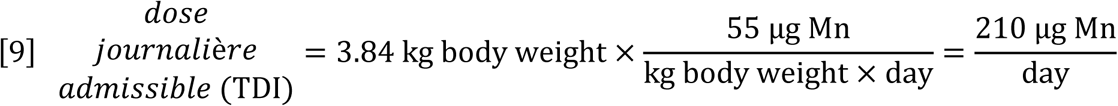

With an intake of 0.619_7…_ L of prepared product per day for a 3-week-old (Table 2), 210 µg Mn/day would be provided by an infant formula with 340 µg Mn/L of product as follows (Equation 10).

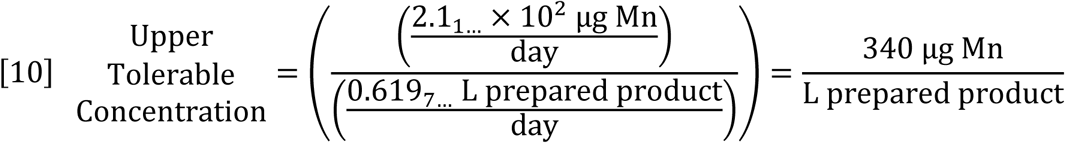

In our recent study of infant formulas and young child nutritional beverages available on the US and French markets, the mean energy content of the products labeled for use by 3-week-olds was 6.6_2…_×10^2^ kcal/L of prepared formula (Table 2) [42]; therefore, 340 µg of Mn/L of prepared product would equal approximately 51 µg of Mn/100 kcal (Equation 11).

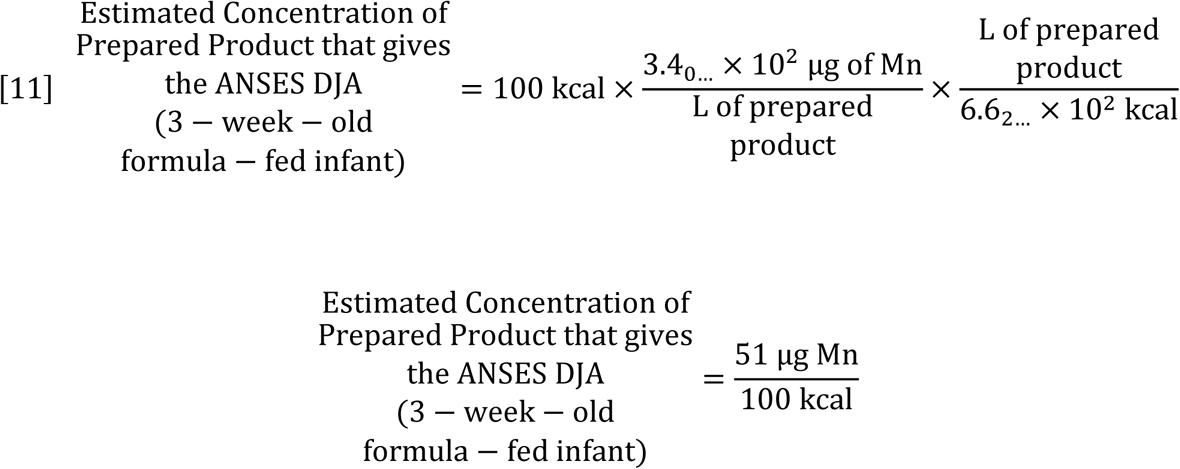

It must be noted that the current infant formula regulations in France stipulate a maximum Mn content of 100 µg Mn/100 kcals [64]. A formula with 100 µg Mn/100 kcals would supply 424 µg Mn/day (Equation 12).

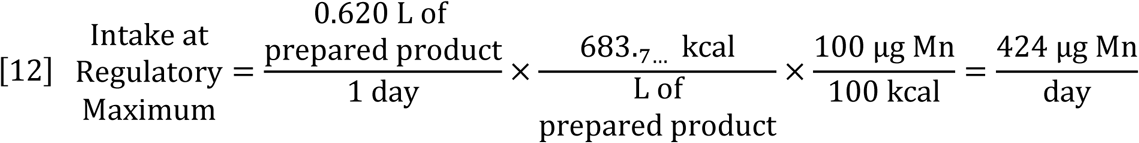

Thus, the current regulatory maximum for infant formula in France exceeds the proposed ANSES *DJA* (TDI) for oral Mn exposure by a factor of 2 (Equations 9, 12) [16,64]. In our market basket survey of products, 2 of the 4 samples from the French market labeled for use by 3-week-olds would lead to exceedances of the ANSES *DJA* (TDI) of 55 μg Mn/kg/day, as would 11 of the 16 samples from the US market labeled as “infant formula” [42].

## Conclusions

### Comparing Mn intakes from breast milk, infant formulas, and young child nutritional beverage products

The concentrations of Mn in human milk at various times after birth reported in the peer-reviewed literature on a volume basis ranged from 0.17 µg/L to 30.27 µg/L, with a weighted mean concentration of 7.7 µg/L (Table 1). Using the regulatory units for infant formulas, this mean concentration for Mn in breast milk would be equivalent to approximately 1.1 µg Mn/100 kcals (Equation 2). Estimated daily intake of Mn/kg of body weight due to breast milk, infant formulas, or young child nutritional beverage products was highest at the youngest age we examined (3 weeks) and declined as age and total body weight increased (Table 2 and Figure 1). Although Mn content of breast milk varies, the overall Mn content and the variability of Mn content of breast milk are much less than that of infant formula products and young child nutritional beverages. The intake of Mn per day from breast milk with the maximum reported concentration of Mn is less than 4 times the daily intake from breast milk with the mean reported concentration of Mn (Tables 1 and 2). In contrast, for infant formulas and young child nutritional beverage products, the daily intake of Mn/kg body weight from the products that we surveyed was between 28 and 520 times the daily intake of Mn/kg body weight from breast milk with the mean reported concentration of Mn (Tables 1 and 2).

### The potential effect of ambient Mn in water on the concentration of manganese in prepared infant formulas and young child nutritional beverages

The estimated total intakes of Mn due to infant formulas and young child nutritional beverages will be substantially higher if reconstituted with water containing a relatively high Mn concentration, but whose Mn concentration is lower than the U.S. EPA Lifetime Health Advisory of 300 µg/L [79] or the WHO Health-Based Value of 400 µg/L [85], e.g. 250 µg/L. If reconstituted with water containing 250 µg Mn/L, some of the products tested from the US market would exceed the U.S. EPA RfD of 140 µg Mn/kg/day (Table 4) [79]. It must be kept in mind that this U.S. EPA RfD was set on the basis of Mn exposures in adults and may not be sufficiently protective of infants and young children, who may be exceptionally sensitive to excess Mn exposures [10,11,89].

### Mn deficiency is highly unlikely

Manganese deficiency is highly unlikely with either exclusive or partial formula-feeding. The IOM has set an Adequate Intake (AI) for 0-6 month-old-infants of 3 µg Mn/day based on an estimated Mn concentration in breast milk of 3.5 µg/L (reference weights not specified) [72]. All products that we tested, both those supplemented with Mn and those not supplemented with Mn, would supply more than the daily AI of Mn for 0-to 6-month-olds (3 µg Mn/day) [72] if used for exclusive feeding [42]. In particular, the unsupplemented infant formula that we tested in our market basket survey that had the minimum Mn concentration (230 µg/L) would supply 29 times the daily Mn intake as breast milk for a 3-week-old and approximately 47 times the AI for Mn for this age stage [42].

### Drinking water guidelines and regulations are inappropriate for dietary exposures

When considering the potential toxicity of nutrient-containing beverages such as infant formulas and young child nutritional beverage products, it is vital that drinking water guidelines or drinking water regulations *not* be used directly to evaluate the safety of the beverage since health-based drinking water guidelines or regulations always incorporate assumptions that are specific to drinking water and inappropriate for dietary exposures (e.g. bioavailability from drinking water, volume of drinking water ingested per day, weight of subjects, and percent allocation from drinking water, food, and other sources). However, health-based drinking water guidelines or regulations typically incorporate a tolerable intake level, based on a NOAEL or LOAEL with units given in the mass of toxic substance or nutrient/kg body weight/day. The tolerable intake levels are used in the *derivations* of the guidelines or regulations, but their values are different from the actual drinking water guidelines or regulations. These differences are due to the mathematical calculations in the guideline or regulation derivations used to account for bioavailability in the substrate, intake volumes, body weight of subjects, percent exposure allocation, and uncertainty factors. Thus, the published tolerable intake levels used in the *derivations* of drinking water guidelines, but not the calculated drinking water-specific guidelines or regulations, could be used to evaluate the potential safety of nutrient-containing beverages or foods such as infant formulas if specific food-based toxicity guidelines or regulations are not available.

### Regulatory tolerable intakes for dietary Mn

While JECFA does not currently have a guideline for Mn in food, in its derivation of a health-based value for Mn in drinking water, the WHO has established a NOAEL of 11 mg Mn/day for adults based on dietary Mn exposures [85,90,91]. This would be equivalent to 180 µg Mn/kg/day using the WHO’s standard assumption of a 60 kg adult (Equation 3). For a 3.84 kg infant (Table 2), this would amount to 700 µg Mn/day (Equation 4), which would be supplied by a formula containing 1,100 µg Mn/100 kcal (Equation 5; Table 2). None of the products that we analyzed in our market basket survey would exceed this level if prepared with water containing 0 µg Mn/L (Table 2) [42].

The IOM declined to establish a UL for dietary Mn intakes for infants, stating “For infants, the UL was judged not determinable because of lack of data on adverse effects in this age group and concern about the infant’s ability to handle excess amounts. To prevent high levels of manganese intake, the only source of intake for infants should be from food or formula” [72]. For young children ages 1-3, the IOM extrapolated by weight the adult UL of 11 mg/day to 2 mg/day (Equation 6). Assuming that young child nutritional beverages provide 21.3% of daily energy intake [57], a proportional UL assigned to nutritional beverages would be 1200 µg Mn/L (Equation 7). None of the young child products from France that we tested in our market basket survey would exceed this level, but one of the toddler formula products from the US would [42]. Beverages labelled to provide complete nutrition would exceed the IOM UL of 2 mg/day if they contain more than 180 µg Mn/100 kcal (Equation 8). One of the medically complete beverages from the US labelled for use by children one year and older would exceed this level [42].

The U.S. EPA has established an RfD of 0.14 mg Mn/kg/day based on dietary intakes. None of the products that we tested in our market basket survey would exceed this intake level if prepared using water with 0 µg/L Mn concentration (Table 2) [42]. However, 2 products from the US labelled for use by infants would exceed this intake level if used to feed 3-week olds and prepared with water with 250 µg Mn/L, a concentration that is relatively high but does not exceed the U.S. EPA HA of 300 µg Mn/L [79] or the WHO HBV of 400 µg Mn/L [85] in drinking water, as would both nutritionally complete medical products labelled for ages one year and older when used to provide all nutritional requirements for 18-month-olds (Table 4) [42].

While the EU does not currently have regulations regarding Mn in dietary exposures, ANSES has recently proposed a maximum tolerable limit for drinking water in France of 60 µg Mn/L based on a tolerable daily intake of 55 µg/kg/day [16,127]. This proposed daily intake would be surpassed by 3-week-old infants drinking formula with greater than 51 µg Mn/100 kcal (Equation 11). It must be noted that current infant formula regulations in France allow a maximum Mn content of 100 µg Mn/100 kcal, which would lead to a daily intake amounting to twice the ANSES proposed maximum daily intake for oral ingestion of Mn (Equation 12) [16,64]. Many products labeled for use by 3-week-olds that we tested from the French and US markets would lead to exceedances of the ANSES tolerable daily intake of 55 µg/kg/day, but some would not, demonstrating that it is possible for infant formulas to be produced with less than 51 µg Mn/100 kcal using current production practices [42].

### Recommendations

Research on neurological effects of Mn exposures in infants and young children is needed to inform risk assessors and regulators. In the meantime, parents and healthcare workers should heed the advice of the WHO, “Breastfeeding is an unequalled way of providing ideal food for the healthy growth and development of infants; it is also an integral part of the reproductive process with important implications for the health of mothers. As a global public health recommendation, infants should be exclusively breastfed for the first six months of life to achieve optimal growth, development and health” [131]. In situations where breast feeding is impossible, the potential for excess Mn intakes causing harm to the brain of the developing infant underscores the need for reducing Mn content of formulas to make it closer to that found in breast milk. Supplementation of infant formulas and young child nutritional beverages with Mn should be discouraged and manufacturers should be urged to reduce Mn content of all formula products to make Mn content more in line with the mean Mn content of breast milk, 1.1 µg Mn/100 kcal (Equation 2). To reach this goal, ingredients with high native Mn content such as soy, rice, or chocolate may need to be limited, with soy- and rice-based formulas used only in case of medical necessity, and optional high Mn-ingredients (eg. chocolate) avoided for infants and young children.

## Acknowledgments

This research did not receive any specific grant from funding agencies in the public, commercial, or not-for-profit sectors. This study received institutional support from Better Life Laboratories, Inc., Norwich University, and the Centre d’Etudes Nucléaires de Bordeaux Gradignan (CENBG). We thank Deborah Ahlers and Tammy Hunt from Norwich University, and Florelle Domart and Francesco Porcaro from CENBG for their valuable assistance with this project.

## ^1^ List of abbreviations

AI: Adequate Intake
ANSES: *Agence nationale de sécurité sanitaire de l’alimentation, de l’environnement et du travail* (French Agency for Food, Environmental and Occupational Health & Safety)
DJA: *dose journalière admissible* (daily admissible dose, tolerable daily intake)
EFSA: European Food Safety Authority
EU: European Union
FAO: Food and Agriculture Organization
HBV: Health-Based Value
INSPQ: *Institut national de santé publique du Québec* (National Institute of Public Health of Québec)
IOM: Institute of Medicine
IQ: intelligence quotient
JECFA: Joint FAO/WHO Expert Committee on Food Additives
LOAEL: Lowest Observed Adverse Effect Level
Mn: Manganese
MnSOD: manganese superoxide dismutase
MR: magnetic resonance
NOAEL: No Observed Adverse Effect Level
PIXE: particle induced X-ray emission
PN: parental nutrition
RBS: Rutherford backscattering
RfD: Reference Dose
SCF: Scientific Committee on Food
T1R: T1 relaxation
TPN: total parenteral nutrition
U.S.: United States Environmental Protection Agency
EPA: Environmental Protection Agency
US: United States

## References

[1] M. Bouchard, F. Laforest, L. Vandelac, D. Bellinger, D. Mergler, Hair manganese and hyperactive behaviors: pilot study of school-age children exposed through tap water, Environ. Health Perspect. 115 (2007) 122–127. https://doi.org/10.1289/ehp.9504.

[2] M.F. Bouchard, S. Sauvé, B. Barbeau, M. Legrand, M.È. Brodeur, T. Bouffard, E. Limoges, D.C. Bellinger, D. Mergler, Intellectual impairment in school-age children exposed to manganese from drinking water, Environ. Health Perspect. 119 (2011) 138–143. https://doi.org/10.1289/ehp.1002321.

[3] S.H. Frisbie, R. Ortega, D.M. Maynard, B. Sarkar B, The concentrations of arsenic and other toxic elements in Bangladesh’s drinking water, Environ. Health Perspect. 110 (2002) 1147–1153. https://doi.org/10.1289/ehp.021101147.

[4] E.J. Mitchell, S.H. Frisbie, B. Sarkar, Exposure to multiple metals from groundwater—a global crisis: Geology, climate change, health effects, testing, and mitigation, Metallomics 3 (2011) 874–908. https://doi.org/10.1039/c1mt00052g.

[5] G.A Wasserman, X. Liu, F. Parvez, H. Ahsan, P. Factor-Litvak, A. van Geen, V. Slavkovich, N.J. Loiacono, D. Levy, Z. Cheng, J.H. Graziano, Water arsenic exposure and children’s intellectual function in Araihazar, Bangladesh, Environ. Health Perspect. 112 (2004) 1329–1333. https://doi.org/10.1289/ehp.9501.

[6] G.A. Wasserman, X. Liu, F. Parvez, H. Ahsan, D. Levy, P. Factor-Litvak, J. Kline, A. van Geen, V. Slavkovich, N.J. Lolacono, Z. Cheng, Y. Zheng, J.H. Graziano, Water manganese exposure and children’s intellectual function in Araihazar, Bangladesh, Environ. Health Perspect. 114 (2006) 124–129. https://doi.org/10.1289/ehp.8030.

[7] L.A. Dion, D. Saint-Amour, S. Sauvé, B. Barbeau, D. Mergler, M.F. Bouchard. Changes in water manganese levels and longitudinal assessment of intellectual function in children exposed through drinking water, Neurotoxicology 64 (2018) 118–125. https://doi.org/10.1016/j.neuro.2017.08.015.

[8] P. Grandjean, P. Landrigan, Neurobehavioural effects of developmental toxicity, Lancet Neurol. 13 (2014) 330–38. https://doi.org/10.1016/S1474-4422(13)70278-3.

[9] S.S. Kullar, K. Shao, C. Surette, G. Foucher, D. Mergler, P. Cormier, D.C. Bellinger, B. Barbeau, S. Sauvé, M.F. Bouchard, 2019, A benchmark concentration analysis for manganese in drinking water and IQ deficits in children, Environ. Int. 130, e104889. https://doi.org/doi:10.1016/j.envint.2019.05.083.

[10] R. Lucchini, D. Placidi, G. Cagna, C. Fedrighi, M. Oppini, M. Peli, S. Zoni, Manganese and developmental neurotoxicity, Adv. Neurobiol. 18 (2017) 13–34. https://doi.org/10.1007/978-3-319-60189-2_2.

[11] J.A. Menezes-Filho, M. Bouchard, P. Sarcinelli, J.C. Moreira, Manganese exposure and the neuro psychological effect on children and adolescents: a review, Rev. Panam. Salud Public 26 (2009) 541–8. https://doi.org/10.1590/s1020-49892009001200010.

[12] A.P. Neal, T.R. Guilarte, Mechanisms of lead and manganese neurotoxicity, Toxicol. Res. (Camb.) 2 (2013) 99–114. https://doi.org/10.1039/C2TX20064C.

[13] T.V. Peres, M.R.C. Schettinger, P. Chen, F. Carvalho, D.S. Avila, A.B. Bowman, 2016. Manganese-induced neurotoxicity: a review of its behavioral consequences and neuroprotective strategies, B.M.C. Pharm. Tox. 17, e57. https://doi.org/10.1186/s40360-016-0099-0.

[14] M. Rodriguez-Barranco, M. Lacasaña, C. Aguilar-Garduño, J. Alguacil, F. Gil, B. González-Alzaga, A. Rojas-García, Association of arsenic, cadmium and manganese exposure with neurodevelopment and behavioural disorders in children: A systematic review and meta-analysis, Sci. Total Environ. 454-455 (2013) 562–577. https://doi.org/10.1016/j.scitotenv.2013.03.047.

[15] S. Zoni, R.G. Lucchini, Manganese exposure: cognitive, motor and behavioral effects on children: a review of recent findings, Curr. Opin. Pediatr. 25 (2013) 255–260. https://doi.org/10.1097/MOP.0b013e32835e906b.

[16] Agence nationale de sécurité sanitaire de l’alimentation, de l’environnement et du travail (French Agency for Food, Environmental and Occupational Health & Safety), Avis de l’Agence nationale de sécurité sanitaire de l’alimentation, de l’environnement et du travail relatif à la détermination d’une valeur sanitaire maximale admissible pour le manganèse dans l’eau destinée à la consommation humaine (Opinion of the National Agency for Food Safety, the Environment and Labor concerning the determination of a maximum allowable health value for manganese in water destined for human consumption. https://www.anses.fr/en/system/files/EAUX2016SA0203.pdf, 2018 (accessed 13 February 2018).

[17] Minnesota Department of Health, Toxicological summary for: Manganese. Health based guidance for water, Minnesota Department of Health, St Paul, MN, 2018.

[18] M. Valcke, M.-H. Bourgault, S. Haddad, M. Bouchard, D. Gauvin, P. Levallois, 2018. Deriving a drinking water guideline for a non-carcinogenic contaminant: The case of manganese, Int. J. of Environ. Res. and Pub. Health 15, e1293. https://doi.org/10.3390/ijerph15061293.

[19] J.L. Pomeranz, M.J. Romo Palafox, J.L. Harris, Toddler drinks, formulas, and milks: Labeling practices and policy implications, Prev. Med. 109 (2018) 11–16. https://doi.org/10.1016/j.ypmed.2018.01.009.

[20] F.M. Al-Awadi, T.S. Srikumar, Trace-element status in milk and plasma of Kuwaiti and non-Kuwaiti lactating mothers, Nutr. 16 (2000) 1069–1073. https://doi.org/10.1016/s0899-9007(00)00426-3.

[21] A.A. Almeida, C.M. Lopes, A.M. Silva, E. Barrado, Trace elements in human milk: Correlation with blood levels, inter-element correlations and changes in concentration during the first month of lactation, J. Trace Elem. Med. Biol. 22 (2008) 196–205. https://doi.org/10.1016/j.jtemb.2008.03.007.

[22] R.R. Anderson, Comparison of trace elements in milk of four species, J. Dairy Sci. 75 (1992) 3050–3055. https://doi.org/10.3168/jds.S0022-0302(92)78068-0.

[23] E. Aquilio, R. Spagnoli, S. Seri, G. Bottone, G. Spennati, Trace element content in human milk during lactation of preterm newborns, Biol Trace Elem Res. 51 (1996) 63–70. https://doi.org/10.1007/BF02790148.

[24] J. Arnaud, A. Favier, Copper, iron, manganese and zinc contents in human colostrum and transitory milk of French women, Sci. Total Environ. 159 (1995) 9–15. https://doi.org/10.1016/0048-9697(94)04314-d.

[25] K.L. Björklund, M. Vahter, B. Palm, M. Grandér, S. Lignell, M. Berglund, 2012. Metals and trace element concentrations in breast milk of first time healthy mothers: A biological monitoring study. Environ. Health. 11, e92. https://doi.org/10.1186/1476-069X-11-92.

[26] B. Bocca, A. Alimonti, L. Giglio, M.D. Pasquale, S. Caroli, A. Ambruzzi, A.P. Bocca, E. Coni, Nutritive significance of element speciation in breast milk, Adv. Exp. Med. Biol. 478 (2000) 385–386. https://doi.org/10.1007/0-306-46830-1_39.

[27] C.E. Casey, K.M. Hambidge, M.C. Neville, Studies in human lactation: zinc, copper, manganese and chromium in human milk in the first month of lactation, Am. J. Clin. Nutr. 41 (1985) 1193–1200. https://doi.org/10.1093/ajcn/41.6.1193.

[28] E. Coni, B. Bocca, B. Galoppi, A. Alimonti, S. Caroli, Identification of chemical species of some trace and minor elements in mature breast milk, Microchem J. 67 (2000) 187–194. https://doi.org/10.1016/S0026-265X(00)00116-8.

[29] K. Dörner, S. Dziadzka, A. Hohn, J. Schaub, Longitudinal manganese and copper balances in young infants and preterm infants fed on breast-milk and adapted cow’s milk formulas, Br J Nutr. 61 (1989) 559–572. https://doi.org/10.1079/bjn19890143.

[30] J.K. Friel, W.L. Andrews, S.E. Jackson, H.P. Longerich, C. Mercer, A. Mcdonald, B. Dawson, B. Sutradhar, Elemental composition of human milk from mothers of premature and full-term infants during the first 3 months of lactation, Biol. Trace Elem. Res. 67 (1999) 225–247. https://doi.org/10.1007/BF02784423.

[31] A.A. Kinsara, S.M. Farid, Concentration of trace elements in human and animal milk in Jeddah, Saudi Arabia, Med. J. Islamic World Acad. Sci. 16 (2006) 181–188.

[32] L.D. Klein, A.A. Breakey, B. Scelza, C. Valeggia, G. Jasienska, K. Hinde, 2017. Concentrations of trace elements in human milk: Comparisons among women in Argentina, Namibia, Poland, and the United States, PLoS ONE. 12, e0183367. https://doi.org/10.1371/journal.pone.0183367.

[33] M. Krachler, T. Prohaska, G. Koellensperger, E. Rossipal, G. Stingeder, Concentrations of selected trace elements in human milk and in infant formulas determined by magnetic sector field inductively coupled plasma–mass spectrometry, Biol. Trace Elem. Res. 76 (2000) 97–112. https://doi.org/10.1385/BTER:76:2:97.

[34] M. Leotsinidis, A. Alexopoulos, E. Kostopoulou-Farri, Toxic and essential trace elements in human milk from Greek lactating women: Association with dietary habits and other factors, Chemosphere 61 (2005) 238–247. https://doi.org/10.1016/j.chemosphere.2005.01.084.

[35] C. Li, N.W. Solomons, M.E. Scott, K.G. Koski, Minerals and trace elements in human breast milk are associated with Guatemalan infant anthropometric outcomes within the first 6 months, J. Nutr. 146 (2016) 2067–2074. https://doi.org/10.3945/jn.116.232223.

[36] R.M. Parr, E.M. Demaeyer, V.G. Iyengar, A.R. Byrne, G.F. Kirkbright, G. Schöch, L. Niinistö, O. Pineda, H.L. Vis, Y. Hofander, A. Omololu, Minor and trace elements in human milk from Guatemala, Hungary, Nigeria, Philippines, Sweden, and Zaire. Biol. Trace Elem. Res. 29 (1991) 51–75. https://doi.org/10.1007/bf03032674.

[37] J. Qian, T. Chen, W. Lu, S. Wu, J. Zhu, Breast milk macro- and micronutrient composition in lactating mothers from suburban and urban Shanghai, J. Paediatr. Child Health. 46 (2010) 115–120. https://doi.org/10.1111/j.1440-1754.2009.01648.x.

[38] E. Rossipal, M. Krachler, Pattern of trace elements in human milk during the course of lactation, Nutr, Res. 18 (1998) 11–24. https://doi.org/10.1016/S0271-5317(97)00196-6.

[39] D. Stastny, R.S. Vogel, M.F. Picciano, Manganese intake and serum manganese concentration of human milk-fed and formula fed infants. Am J Clin Nutr. 39 (1984) 872–878. https://doi.org/0.1093/ajcn/39.6.872.

[40] N. Yamawaki, M. Yamada, T. Kan-no, T. Kojima, T. Kaneko, A. Yonekubo, Macronutrient, mineral and trace element composition of breast milk from Japanese women, J. Trace Elem. Med. Biol. 19 (2005) 171–181. https://doi.org/10.1016/j.jtemb.2005.05.001.

[41] R. Rosenthal, M.R. DiMatteo, Meta-analysis: Recent developments in quantitative methods for literature reviews, Annu. Rev. Psychol. 52 (2001) 59–82. https://doi.org/10.1146/annurev.psych.52.1.59.

[42] S.H. Frisbie, E.J. Mitchell, A. Carmona, S. Roudeau, F. Domart, R. Ortega, 2019. Manganese levels in infant formula and young child nutritional beverages in the United States and France: Comparison to breast milk and regulations. PLoS ONE. 14, e0223636. https://doi.org/10.1371/journal.pone.0223636.

[43] C. Teddlie, F. Yu, Mixed methods sampling, A typology with examples, J Mix. Methods Res. 1 (2007) 77–100. https://doi.org/10.1177/1558689806292430.

[44] N.H.M. Taufek, Essential trace element analysis of human breast milk. A thesis submitted for the degree of Doctor of Philosophy at The University of Queensland School of Pharmacy. University of Queensland, Queensland, Australia. 2016, p. 105.

[45] E. Vuori, A longitudinal study of manganese in human milk, Acta.Paediatr. Scand. 68 (1979) 571–573. https://10.1111/j.1651-2227.1979.tb05057.x.

[46] C.M. Barrera, H.C. Hamner, C.G. Perrine, K.S. Scanlon, Timing of introduction of complementary foods to US infants, National Health and Nutrition Examination Survey 2009-2014, J. Acad. Nutr. Diet. 118 (2018) 464–470. https://doi.org/10.1016/j.jand.2017.10.020.

[47] K. Synnott, J. Bogue, C.A. Edwards, J.A. Scott, S. Higgins, E. Norin, D. Frias, S. Amarri, R. Adam, Parental perceptions of feeding practices in five European countries: An exploratory study, Eur. J. Clin. Nutr. 61 (2007) 946–956. https://doi.org/10.1038/sj.ejcn.1602604.

[48] K.G. Dewey, M.J. Heinig, L.A. Nommsen, J.M. Peerson, B. Lönnerdal, Growth of breast-fed and formula-fed infants from 0 to 18 months: The DARLING study. Pediatr. 89 (1992) 1035–1041. https://doi.org/10.1136/adc.81.5.395.

[49] E. Sievers, H.D. Oldigs, R. Santer, J. Schaub, Feeding patterns in breast-fed and formula-fed infants, Ann Nutr Metab. 46 (2002) 243–248. https://doi.org/10.1159/000066498.

[50] D. Stastny, R.S. Vogel, M.F. Picciano, Manganese intake and serum manganese concentration of human milk-fed and formula fed infants, Am. J. Clin. Nutr. 39 (1984) 872–878. https://doi.org/10.1093/ajcn/39.6.872.

[51] K.G. Dewey, R.J. Cohen, L.A. Nommsen-Rivers, M.J. Heinig, Implementation of the WHO Multicentre Growth Reference Study in the United States, Food and Nutr. Bull. 25 (2004) S84–S89. https://doi.org/10.1177/15648265040251s112.

[52] World Health Organization, Child growth standards. https://www.who.int/childgrowth/standards/weight_for_age/en/, 2006 (accessed 16 January 2020).

[53] R.J. Kuczmarski, C.L. Ogden, S.S. Guo, L.M. Grummer-Strawn, K.M. Flegal, Z. Mei, R. Wei, L.R. Curtin, A.F. Roche, C.L. Johnson, 2000 CDC growth charts for the United States: Methods and development, Vital Health Stat. 11 (2002) 1–190.

[54] M. de Onis, C. Garza, A.W. Onyango, E. Borghi, Comparison of the WHO Child Growth Standards and the CDC 2000 Growth Charts, J. Nutr. 137 (2007) 144–148. https://doi.org/10.1093/jn/137.1.144.

[55] P. Scherdel, J. Botton, M.-F. Rolland-Cachera, J. Léger, F. Pelé, P.Y. Ancel, C. Simon, K. Castetbon, B. Salanave, H. Thibault, S. Lioret, S. Péneau, G. Gusto, M.A. Charles, B. Heude, 2015, Should the WHO growth charts be used in France?. PLoS One. 10, e0120806. https://doi.org/10.1371/journal.pone.0120806.

[56] I.G. Macey, The composition of milks: a compilation of the comparative composition and properties of human, cow, and goat milk, colostrum, and transitional milk, National Academy of Sciences, National Research Council, Washington D.C., 1953, p. 62.

[57] M. Fantino, E. Gourmet, *Apports nutritionnels en France en 2005 chez les enfants non allaités âgés de moins de 36 mois* (Nutrient intakes in 2005 by non-breast fed French children of less than 36 months), Archives de Pédiatrie 15 (2008) 446–455. https://10.1016/j.arcped.2008.03.002.

[58] W.M. Haynes, (Ed.), CRC handbook of chemistry and physics, ninety-third ed., CRC Press, Boca Raton, Florida, 2012, p. 749.

[59] D. Mandel, R. Lubetzky, S. Dollberg, S. Barak, F.B. Mimouni, Fat and energy contents of expressed human breast milk in prolonged lactation, Pediatr. 116 (2005) e432–e435. https://doi.org/10.1542/peds.2005-0313.

[60] S. Wünschmann, I. Kühn, H. Heidenreich, S. Fränzle, O. Wappelhorst, B. Markert, Transfer von elementen in die muttermilch (Transfer of elements into mother’s milk), Internationales Hochschulinstitut Zittau, Zittau, Germany, 2003, p. 47–49.

[61] Code of Federal Regulations (21 CFR 107.100), Title 21 - Food and drugs. Part 107 - Infant formula. Subpart D - Nutrient requirements. §107.100 - Nutrient specifications. https://www.ecfr.gov/cgi-bin/text-idx?SID=7e29e474204f59ab0de67762bb5991cb&mc=true&node=se21.2.107_1100&rgn=div8, 2017 (accessed 17 November 2017).

[62] Codex Alimentarius Commission, Standard for infant formula and formulas for special medical purposes intended for infants. CODEX STAN 72-1981. http://www.fao.org/fao-who-codexalimentarius/standards/list-of-standards/en/, 2016. (accessed 9 November 9 2017) [link no longer functional 18 November 2019].

[63] European Commission, Official Journal of the European Union. Commission delegated regulation (EU) 2016/127 of 25 September 2015 supplementing Regulation (EU) No 609/2013 of the European Parliament and of the Council as regards the specific compositional and information requirements for infant formula and follow-on formula and as regards requirements on information relating to infant and young child feeding. https://eur-lex.europa.eu/legal-content/EN/TXT/?uri=CELEX%3A32016R0127, 2015 (accessed 1 February 2016).

[64] ECEC0771649A, Journal Officiel de la République Française (Official Journal of the French Republic). Arrêté du 11 avril 2008 relatif aux préparations pour nourrissons et aux préparations de suite et modifiant l’arrêté du 20 septembre 2000 relatif aux aliments diététiques destinés à des fins médicales spéciales (Order of 11 April 2008 on infant formulas and follow-on formulas and amending the order of 20 September 2000 on dietary foods for special medical purposes). n°0096 du 23 avril 2008 page 6700, texte n°18, Version consolidée au 15 janvier (n°0096 of April 23, 008 page 6700, text n°18, consolidated version on January 15, 2018). https://www.legifrance.gouv.fr/affichTexte.do?cidTexte=JORFTEXT000018685743&dateTexte=20180115, 2018 (accessed 15 January 2018).

[65] L.A. Nommsen, C.A. Lovelady, M.J. Heinig, B. Lönnerdal, K.G. Dewey, Determinants of energy, protein, lipid, and lactose concentrations in human milk during the first 12 mo of lactation: The DARLING study, Am. J. Clin. Nutr. 53 (1991) 457–465. https://doi.org/10.1093/ajcn/53.2.457.

[66] B. Koletzko, S. Baker, G. Cleghorn, U.F. Neto, S. Gopalan, O. Hernell, Q.S. Hock, P. Jirapinyo, B. Lonnerdal, P. Pencharz, H. Pzyrembel, J. Ramirez-Mayans, R. Shamir, D. Turck, Y. Yamashiro, D. Zong-Yi, Global standard for the composition of infant formula: Recommendations of an ESPGHAN coordinated international expert group, J. Pediatr. Gastroenterol. Nutr. 41 (2005) 584–599. https://doi.org/10.1097/01.mpg.0000187817.38836.42.

[67] United States Environmental Protection Agency, Child-specific exposure factors handbook, EPA/600/R-06/096F. https://ofmpub.epa.gov/eims/eimscomm.getfile?p_download_id=484738, 2008 (accessed 22 May 2018).

[68] Institut national de santé publique du Québec, Lignes directrices pour la realization des évaluations du risque toxicologique d’origine environnementale au Québec (Guidelines for the Conduct of Toxicological Risk Assessments of Environmental Origin in Quebec). https://www.inspq.qc.ca/pdf/publications/1440_LignesDirectRealEvaRisqueToxicoOrigEnviroSanteHum.pdf, 2012 (accessed 14 February, 2019).

[69] Codex Alimentarius Commission, Standard for infant formula and formulas for special medical purposes intended for infants. CODEX STAN 72-1981. http://www.fao.org/fao-who-codexalimentarius/sh-proxy/fr/?lnk=1&url=https://workspace.fao.org/sites/codex/Standards/CODEx+STAN+72-1981/CXS_072e.pdf, 2007 (accessed 17 March 2018).

[70] European Food Safety Authority, 2014. Scientific opinion on the essential composition of infant and follow-on formulae. EFSA J. 12, e3760. https://doi.org/10.2903/j.efsa.2014.3760.

[71] Agence française de sécurité sanitaire des aliments (French Food Safety Agency), Avis de l’Agence française de sécurité sanitaire des aliments relatif à l’évaluation d’un aliment diététique destiné aux nourrissons, enrichi en manganèse (Opinion of the French Food Safety Agency concerning the evaluation of a dietary food for infants, enriched in manganese). https://www.anses.fr/fr/system/files/NUT2001sa0090.pdf, 2002 (accessed 17 March 2018).

[72] Institute of Medicine, Dietary Reference Intakes for vitamin A, vitamin K, arsenic, boron, chromium, copper, iodine, iron, manganese, molybdenum, nickel, silicon, vanadium, and zinc, National Academy Press, Washington, D.C., 2001, pp. 412–414.

[73] World Health Organization, The quantity and quality of breast milk, World Health Organization: Geneva, 1985, pp. 31–51.

[74] M.C. Kay, E.B. Welker, E.E. Jacquier, M.T. Story, 2018. Beverage consumption patterns among infants and young children (0–47.9 months): Data from the Feeding Infants and Toddlers Study, 2016, Nutrients. 10, e825. https://doi.org/10.3390/nu10070825.

[75] B. Lönnerdal, Regulation of mineral and trace elements in human milk: Exogenous and endogenous factors, Nutr Rev. 58 (2000) 223–229. https://10.1111/j.1753-4887.2000.tb01869.x.

[76] Agencenationale de sécurité sanitaire de l’alimentation, de l’environnement et du travail; French Agency for Food, Environmental and Occupational Health & Safety. (ANSES). AVIS de l’Agence nationale de sécurité sanitaire de l’alimentation, de l’environnement et du travail relatif à la détermination d’une valeur sanitaire maximale admissible pour le manganèse dans l’eau destinée à la consommation humaine. (Opinion of the National Agency for Food Safety, the Environment and Labor concerning the determination of a maximum admissible health value for manganese in water destined for human consumption) 2018. https://www.anses.fr/fr/system/files/EAUX2016SA0203.pdf. Accessed February 26, 2019.

[77] Centers for Disease Control and Prevention, Infant feeding. https://www.cdc.gov/breastfeeding/pdf/ifps/data/ifps2_tables_ch3.pdf, 2014 (accessed 6 October 2017).

[78] European Economic Community, Synthesis report on the quality of drinking water in the member states of the European Union in the period 1999-2001, Directive 80/778/EEC. https://circabc.europa.eu/sd/a/58220131-ecc4-49f9-9ad1-bbb6e9e79578/report1999-2001.pdf, 2008 (accessed 1 February 2018).

[79] United States Environmental Protection Agency, Drinking water health advisory for manganese, EPA-822-R-04-003. https://www.epa.gov/sites/production/files/2014-09/documents/support_cc1_magnese_dwreport_0.pdf, 2004 (Accessed 31 January 2018).

[80] S.H. Frisbie, E.J. Mitchell, H. Dustin, D.M. Maynard, B. Sarkar B, World Health Organization discontinues its drinking-water guideline for manganese, Environ. Health Perspect. 120 (2012) 775–778. https://doi.org/10.1289/ehp.1104693.

[81] F. Malard, J.L. Reygrobellet, R. Laurent, Spatial distribution of hypogean invertebrates in an alluvial aquifer polluted by iron and manganese, Rhône River, France, Verh Internat Verein Limnol. 26 (1998) 1590–1594. https://doi.org/10.1080/03680770.1995.11900996.

[82] P. Stackelberg, Groundwater quality in the glacial aquifer system, United States, U.S. Geological Survey Fact Sheet 2017–3055. https://pubs.er.usgs.gov/publication/fs20173055, 2017 (accessed 31 January 2018). doi: https://doi.org/10.3133/fs20173055.

[83] Town of Randolph, Vermont. Drinking water at the Randolph Village water system contains elevated levels of manganese from the Pearl Street well. http://randolphvt.org/vertical/sites/%7BD7EA543D-4DEE-41D3-BD57-86E65A3C936B%7D/uploads/Manganese_2015_PUBLIC_NOTICE_102915.pdf, 2015 (Accessed 31 January 2018). [link no longer functional 18 November 2019].

[84] SANPO720201A. Journal Officiel de la République Française. (Official Journal of the French Republic). Arrêté du 11 janvier 2007 relatif aux limites et références de qualité des eaux brutes et des eaux destinées à la consommation humaine mentionnées aux articles R. 1321-2, R. 1321-3, R. 1321-7 et R. 1321-38 du code de la santé publique. (Order of 11 January 2007 on the quality limits and references for raw water and water intended for human consumption mentioned in Articles R. 1321-2, R. 1321-3, R. 1321-7 et R. 1321-38 of the code of public health). 6 February, 2007, text 17 of 121. https://beta.legifrance.gouv.fr/loda/texte_lc/LEGITEXT000031642768/ 2007, (accessed 12 February 2018).

[85] World Health Organization, Guidelines for drinking-water quality. Fourth edition incorporating the first addendum, fourth ed, World Health Organization, Geneva, 2017, pp.386–387.

[86] National Research Council, Recommended dietary allowances, tenth ed. https://www.nap.edu/download/1349, 1989 (accessed 13 March 2018).

[87] H.A. Schroeder, J.J. Balassa, I.H. Tipton. Essential trace elements in man: Manganese. A Study in homeostasis. J. Chron. Dis. 19 (1966) 545–571. https://doi.org/10.1016/0021-9681(66)90094-4.

[88] World Health Organization, Trace elements in human nutrition. apps.who.int/iris/bitstream/10665/37931/2/9241561734_eng.pdf, 1973 (accessed 13 March 2017).

[89] P. Chen, S. Chakraborty, T.V. Peres, A.B. Bowman, M. Aschner. Manganese-induced neurotoxicity: From *C. elegans* to humans, Toxicol. Res. 4 (2015) 191–202. https://doi.org/10.1039/C4TX00127C.

[90] World Health Organization, Guidelines for drinking-water quality, third ed, World Health Organization, Geneva, 2004, pp. 397–399, 492.

[91] World Health Organization, Manganese in drinking-water: Background document for development of WHO guidelines for drinking-water quality. http://apps.who.int/iris/handle/10665/75376, 2004. (accessed 9 October 2017).

[92] World Health Organization, Guidelines for drinking-water quality, fourth ed, World Health Organization, Geneva, 2011, pp. 386–387, 471.

[93] World Health Organization, Manganese in drinking-water: Background document for development of WHO guidelines for drinking-water quality. http://www.who.int/entity/water_sanitation_health/dwq/chemicals/manganese.pdf, 2011. (accessed 9 June 2014).

[94] F. H. Nielsen, Manganese, molybdenum, boron, chromium, and other trace elements, in: J.W. Erdman, Jr., I.A. Macdonald, S.H. Zeisel, (Eds.), Present Knowledge in Nutrition, tenth ed, Wiley-Blackwell, Ames, IA, 2012, pp. 586–607.

[95] S.F. Velazquez, J.T. Du, Derivation of the reference dose for manganese, in: W. Mertz, C.O. Abernathy, S.S. Olin (Eds.), Risk Assessment of Essential Elements. ISLI Press, Washington, DC, 1994, pp. 253–266.

[96] L. Davidsson, Å. Cederblad, B. Lönnerdal, B. Sandström, Manganese absorption from human milk, cow’s milk and infant formulas, in: L.S. Hurley, C.L. Keen, B. Lönnerdal, R.B. Rucker (Eds.) Trace Elements in Man and Animals 6, Plenum Press, New York, 1988, pp. 511–512.

[97] J.L. Aschner, M. Aschner, Nutritional aspects of manganese homeostasis, Mol. Aspects Med. 26 (2005) 353–362. https://doi.org/10.1016/j.mam.2005.07.003.

[98] K. Ljung, M. Vahter, Time to re-evaluate the guideline value for manganese in drinking water?, Environ. Health Persp. 115 (2007) 1533–1538. https://doi.org/10.1289/ehp.10316.

[99] C.D. Davis, L. Zech, J.L. Gregger, Manganese metabolism in rats: an improved methodology for assessing gut endogenous losses, Proc. Soc. Exper. Biol. Med. 202 (1993) 103–108. https://doi.org/10.3181/00379727-202-43518.

[100] B. Sandström, L. Davidsson, R. Eriksson, M. Alpsten, C. Bogentoft. Retention of selenium (75Se), Zinc (65Zn) and manganese (54Mn) in humans after intake of a labelled vitamin and mineral supplement, J. Trace Elem. Elect. H. 1 (1987) 33–8.

[101] T.T. Tran, W. Chowanadisai, B. Lönnerdal, L. Le, M. Parker, A. Chicz-Demet, F. M. Crinella, Effects of neonatal dietary manganese exposure on brain dopamine levels and neurocognitive functions, Neurotoxicology 23 (2002) 645–651. https://doi.org/10.1016/s0161-813x(02)00068-2.

[102] S.M. Lasley, C.A. Fornal, S. Mandal, B.J. Strupp, S.A. Beaudin, D.R. Smith, Early postnatal manganese exposure reduces rat cortical and striatal biogenic amine activity in adulthood. Toxicol Sci. 173 (2020)144-155. https://doi.org/toxsci/kfz208.

[103] H. Gong, T. Amemiya, Optic nerve changes in manganese-deficient rats. Exp Eye Res. 68 (1999) 313–320. https://doi.org/10.1006/exer.1998.0602.

[104] M.B. Richards, Manganese in relation to nutrition, Biochem. J. 24 (1930) 1572–1590. https://doi.org/10.1042/bj0241572.

[105] P.J. Schaible, S.L. Bandemer, J.A. Davidson, Michigan Technical Bulletin 159: The Manganese Content of Feedstuffs and its Relation to Poultry Nutrition, Michigan State College Agricultural Experiment Station, East Lansing, MI, 1938. pp. 6–23.

[106] L. Strause, P. Saltman, J. Glowacki, The effect of deficiencies of manganese and copper on osteoinduction and on resorption of bone particles in rats, Calcif. Tissue Int. 41 (1987) 145–150. https://doi.org/10.1007/BF02563794.

[107] E.J. Underwood, Trace Elements in Human and Animal Nutrition, third ed, Academic Press, Cambridge, MA, 1971, pp. 177–207.

[108] M.A. de la Fuente, A. Olano, M. Juârez. Distribution of calcium, magnesium, phosphorus, zinc, manganese, copper and iron between the soluble and colloidal phases of ewe’s and goat’s milk, Lait, 77 (1997) 515–520. https://doi.org/10.1051/lait:1997437.

[109] J. A. Milner, Trace minerals in the nutrition of children, J. Pediatr. 117 (1990) S147–S155. https://doi.org/10.1016/s0022-3476(05)80013-7.

[110] B. Lönnerdal, Nutritional aspects of soy formula, Acta Paediatr. Suppl. 402 (1994) 105–108. https://doi.org/10.1111/j.1651-2227.1994.tb13371.x.

[111] S. Bagdat, E.K. Baran, F. Tokay, Element fractionation analysis for infant formula and food additives by inductively coupled plasma optical emission spectrometry, Int. J. Food Sci. Tech. 49 ((2014) 392–398. https://doi.org/10.1111/ijfs.12312.

[112] R.S. Gibson, C.A. Scythes, Trace element intakes of women, Br. J. Nutr. 48 (1982) 241–248. https://doi.org/10.1079/BJN19820110.

[113] N.L. Kent, R.A. McCance, The absorption and excretion of ‘minor’ elements by man. 2. Cobalt, nickel, tin and manganese, Biochem. J. 35 (1941) 877–883. https://10.1042/bj0350877.

[114] S.D. Soman, V.K. Panday, K.T. Joseph, S.J. Raut, Daily intake of some major and trace elements, Health Phys. 17 (1969) 35–40. https://10.1097/00004032-196907000-00004.

[115] J.K. Kresovich, C.M. Bulka, B.T. Joyce, P.S. Vokonas, J. Schwartz, A.A. Baccarelli, E.A. Hibler, L. Hou, The inflammatory potential of dietary manganese in a cohort of elderly men, Biol. Trace Element Res. 183 (2018) 183:49-57. https://doi.org/10.1007/s12011-017-1127-7.

[116] R.G. Banta, W.R. Markesbery, Elevated manganese levels associated with dementia and extrapyramidal signs, Neurology 27 (1977) 213–216. https://doi.org/10.1212/wnl.27.3.213.

[117] M. Holzgraefe, W. Poser, H. Kijewski, W. Beuche, Chronic enteral poisoning caused by potassium permanganate: a case report, J. Toxicol. Clin. Toxicol. 24 (1986) 235–244. https://doi.org/10.3109/15563658608990461.

[118] M.J. Schuh, Possible Parkinson’s Disease induced by chronic manganese supplement ingestion, Consult. Pharm. 31 (2016) 698–703. https://doi.org/10.4140/TCP.n.2016.698.

[119] J.L. Aschner, A. Anderson, J.C. Slaughter, M. Aschner, S. Steele, A. Beller, A. Mouvery, H.M. Furlong, N.L. Maitre, Neuroimaging identifies increased manganese deposition in infants receiving parenteral nutrition, Am. J. Clin. Nutr. 102 (2015) 1482–1489. https://doi.org/10.3945/ajcn.115.116285.

[120] P. Brna, K. Gordon, J.M. Dooley, V. Price, Manganese toxicity in a child with iron deficiency and polycythemia, J. Child Neurol. 26 (2011) 891–894. https://doi.org/10.1177/0883073810393962.

[121] A. Woolf, R. Wright, C. Amarasiriwardena, D. Bellinger, A child with chronic manganese exposure from drinking water, Environ. Health Perspect. 110 (2002) 613–616. https://doi.org/10.1289/ehp.02110613.

[122] J.L. Greger, Nutrition versus toxicity of manganese in humans: Evaluation of potential biomarkers, Neurotoxicology 20 (1999) 205–212.

[123] W. Laohaudomchok, X. Lin, R.F. Herrick, S.C. Fang, J.M. Cavallari, D.C. Christiani, M.G. Weisskopf, Toenail, blood and urine as biomarkers of manganese exposure, J. Occup. Environ. Med. 53 (2011) 506–510. https://doi.org/10.1097/JOM.0b013e31821854da.

[124] W. Zheng, S.X. Fu, U. Dydak, D.M. Cowan, Biomarkers of manganese intoxication, Neurotoxicology 32 (2011) 1–8. https://doi.org/10.1016/j.neuro.2010.10.002.

[125] Codex Alimentarius Commission, General standard for contaminants and toxins in food and feed. CODEX STAN 193-1995. http://www.fao.org/input/download/standards/17/CXS_193e_2015.pdf, 2015 (accessed 5 March 2018).

[126] Agency for Toxic Substances and Disease Registry, Toxicological profile for manganese. https://www.atsdr.cdc.gov/toxprofiles/tp151.pdf, 2012 (accessed 27 February, 2019).

[127] European Food Safety Authority, Tolerable upper intake levels for vitamins and minerals. https://www.efsa.europa.eu/sites/default/files/efsa_rep/blobserver_assets/ndatolerableuil.pdf, 2006 (accessed 24 October 2017).

[128] Scientific Committee on Food, Opinion of the Scientific Committee on Food on the tolerable upper intake level of manganese. https://ec.europa.eu/food/sites/food/files/safety/docs/sci-com_scf_out80f_en.pdf, 2000 (accessed 13 March 2018).

[129] European Commission, Official Journal of the European Communities 5.12.98. Council directive 98/83/EC of 3 November 1998 on the quality of water intended for human consumption. https://ec.europa.eu/jrc/communities/file/666/download?token=THlHFbU2, 1998. (accessed 20 March 2018).

[130] République Française (Republic of France). Décret n° 2001-1220 du 20/12/01 Relatif aux eaux destinées à la consommation humaine, à l’exclusion des eaux minérales naturelles (Decree No. 2001-1220 of 20/12/01 Relating to water intended for human consumption, excluding natural mineral waters). https://aida.ineris.fr/consultation_document/2955, 2001 (accessed 9 March 2018).

[131] World Health Organization, Global Strategy for Infant and Young Child Feeding, World Health Organization, Geneva, 2003, pp. 7,8.

